# Effective control of treadmilling filaments by plus-end capping

**DOI:** 10.64898/2026.11.22.768371

**Authors:** Xing Ye, Felix Wodaczek, Anđela Šarić, Martin Loose

## Abstract

Bacterial cell division depends on the tubulin homolog FtsZ, which assembles into a membrane-bound cytokinetic Z-ring. FtsZ filaments treadmill, growing at one end and shrinking at the other, and many regulatory proteins control where and when the Z-ring forms. Whether treadmilling itself confers a regulatory advantage, and how inhibitors exploit it, has remained unclear. Here we show that the *Escherichia coli* nucleoid occlusion factor SlmA caps the growing ends of treadmilling FtsZ filaments, converting steady-state subunit turnover into net filament shrinkage. Using *in vitro* reconstitution of membranebound FtsZ filaments with quantitative fluorescence and single-molecule imaging, we find that SlmA shortens filaments and reduces polymer mass at concentrations far below that of FtsZ, behaving like the bona fide capper FtsZ L272E. Structural prediction and mutagenesis identify a previously unrecognized interface between SlmA and the polymerization-competent surface of FtsZ that is required for activity *in vitro* and *in vivo* and that acts together with the known FtsZ C-terminal peptide contact. A minimal stochastic model shows why end capping outperforms sequestration and severing, establishing kinetic asymmetry as a general point of control for treadmilling cytoskeletal polymers.

## 1 Introduction

### 1.1 Prokaryotic cytokinesis is under the control of the Z-ring

In bacteria and many archaea, cytokinesis is governed by the tubulin-like GTPase FtsZ, which assembles into a membrane-associated, cytoskeletal structure known as the Z-ring [1, 2, 3]. FtsZ polymers are highly dynamic and form treadmilling filaments that continuously grow at one end while shrinking at the other [4, 5, 6, 7, 8]. The close coupling between cell division and DNA segregation during cytokinesis is essential for a faithful transmission of genetic information from the mother to their daughter cells [9, 10, 11]. Accordingly, polymerization of FtsZ filaments and Z-ring assembly must be under tight biochemical control. In *Escherichia coli* (*E. coli*) and many other gamma- and beta-proteobacteria, two regulatory systems control where the Z-ring assembles [11, 12]. First, the Min system establishes an inhibitory regulatory gradient to prevent Z-ring assembly at the cell poles [13, 14, 15]. In addition, DNA-binding proteins, the nucleoid-associated protein SlmA (Synthetic Lethal with MinA) in case of *E. coli*, prevent FtsZ polymerization near the dividing chromosomes [16, 17]. Loss of either system causes little disruption to *Escherichia coli* cell division, producing the minicell phenotype in the case of MinCDE or leading to mild cell elongation or slight misplacement of FtsZ rings without SlmA [18, 19]. The absence of both, however, leads to misplaced septa, chromosome guillotining, and anucleate minicell formation and becomes lethal [17, 18, 20].

While the cellular function of Z-ring regulators is known, the underlying molecular mechanisms by which they inhibit polymerization of FtsZ is not well established. So far, mechanistic studies using purified negative regulators of FtsZ polymerization mainly focused on whether they affect polymer mass in sedimentation assays or influence the GTPase activity of FtsZ [18, 21, 22, 23], which helped to identify FtsZ’s conserved C-terminal peptide (CTP) as an important interaction motif. Mutational, genetic and structural biology experiments showed that the CTP interacts with MinC [24, 25] and SlmA [26, 27] possibly competing with membrane anchors FtsA and ZipA that bind to the same motif. Next to the CTP, the globular domain of FtsZ has also been recognized to be an important interaction site for FtsZ regulators. For example, biochemical and structural studies on FtsZ in complex with SulA [28] and the N-terminal domain of MinC [29, 30, 31, 32] as well with MciZ from *Bacillus subtilis* [21] and MipZ from *Caulobacter crescentus* [33] all show that they bind to the bottom surface of FtsZ and therefore share the interaction interface with the incoming FtsZ monomer during filament growth. This mode of binding could have important functional consequences as these proteins would cap the growing end of the treadmilling filament and therefore block further growth. The functional consequences of this interaction for FtsZ polymerization dynamics have not been characterized yet.

Treadmilling is a non-equilibrium process that preserves the steady-state filament length and structural integrity of filaments while continuous subunit loss and addition enables dynamic cytoskeletal remodeling (**fig**. 1**a**). Studies on actin filaments demonstrate that treadmilling provides a powerful regulatory principle: filament-binding proteins can tune subunit on- and off-rates, nucleate new filaments, or block elongation, thereby reshaping filament length, organization, and treadmilling velocity in response to developmental and environmental cues [34, 35, 36]. By suppressing elongation while permitting continued depolymerization, filament capping proteins could in principle result in progressive reductions in FtsZ filament mass [37, 38]. Despite studies that showed that regulators exploit monomer turnover [39], and shorten FtsZ filaments [16, 40, 41], if and how they use treadmilling for regulation has so far not been described and a direct experimental demonstration for how regulatory proteins modulate steady-state filament mass, length and treadmilling velocity has to our knowledge not been achieved yet.

**Figure 1:**
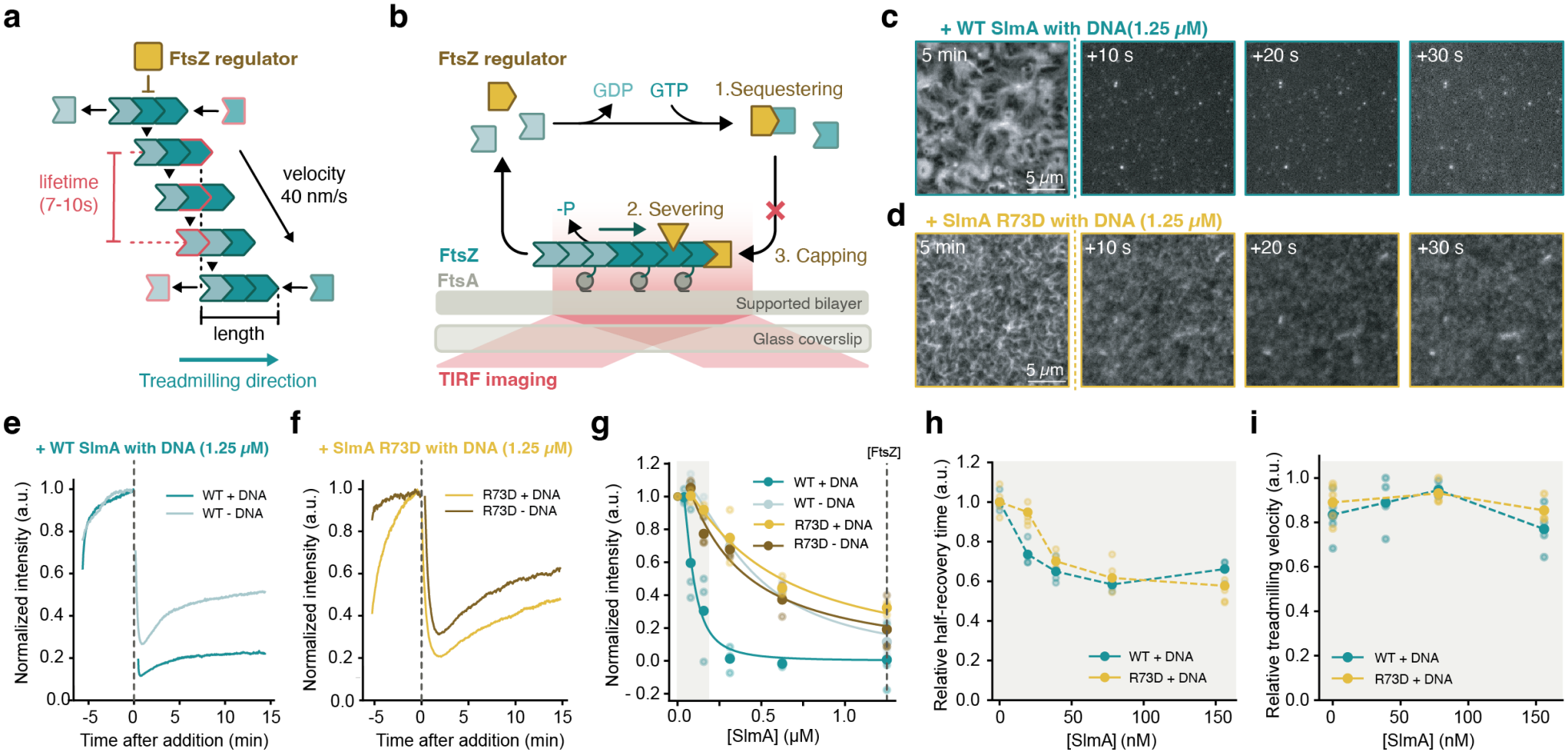
SlmA depolymerizes FtsZ filaments at substoichiometric concentrations. **a)** FtsZ forms treadmilling filaments which can be quantitatively described by their treadmilling velocity, the filament length and their monomer turnover. **b)** Schematic of experimental used and of the different mechanisms of FtsZ regulation, i.e. monomer sequestering, filament capping or severing. Adding **c)** SBS-WT SlmA or **d)** SBS-SlmA R73D (1.25 µM) to membrane-bound patterns of treadmilling FtsZ filaments (1.25 µM) and FtsA (0.4 µM) rapidly removes filaments. **e)** and **f)** Fluorescence intensity curves corresponding to micrographs shown in c) and d), respectively. Curves are representative of *n* = 3 independent experiments per condition. **g)** Decrease of FtsZ fluorescence intensity (**sec. S3.3**) is SlmA concentration dependent. Curves correspond to fitted inhibitory Hill functions. **h)** Addition of SlmA decreases half-recovery time and increases filament turnover before reaching a plateau. At the same time, the treadmilling velocity **i)** stays constant. In g)–i), each dot represents one independent experiment (*n* = 3 per condition).

Here we ask how SlmA controls FtsZ polymerization, and what its mechanism reveals about the regulation of treadmilling filaments in general. SlmA is related to the transcriptional regulator TetR [18] and forms dimers via its helix-turn-helix domain [16]. It binds 24 so-called SlmA-binding sites (SBSs), specific DNA sequences distributed across the chromosome except in the region near the terminus of replication [17]. SBS-bound SlmA antagonizes Z-ring assembly and FtsZ polymerization *in vitro* in a process that relies on filament turnover but does not increase the GTPase activity [17], leading to the proposal of a two-pronged mechanism [25, 42, 43] in which SlmA engages both FtsZ’s CTP [17, 20, 27] and, through a yet unidentified mode of interaction, its globular domain [16, 17, 26, 44]. Direct evidence for this mechanism, and for how it would affect filament length, treadmilling velocity and filament organization, is still missing. Combining *in vitro* reconstitution of membrane-bound, treadmilling FtsZ filaments with quantitative imaging, structural analysis and stochastic modeling, we find that SlmA caps growing filament ends. In a treadmilling polymer, growth and shrinkage occur at opposite ends. Blocking the growing end therefore converts steady-state subunit turnover into net disassembly. Such an effect is not possible for polymers at equilibrium, and it may explain why structurally unrelated FtsZ regulators from different bacteria target the same surface [28, 21, 29, 33].

## 2 Results

### 2.1 SlmA shortens treadmilling filaments even without CTP binding

Binding of SlmA to FtsZ’s C-terminal peptide has been considered to be the primary mechanism to inhibit FtsZ polymerization. Mutations within the CTP of *E. coli* FtsZ (L378 and I374) were found to abrogate the SlmA-FtsZ interaction *in vitro* [26] and a genetic screen in *E. coli* revealed residues involved in SlmA’s antagonism of FtsZ polymerization [20]. These were found to cluster on two adjacent alpha-helices (Phe65, Arg73 on α4 and Leu94, Gly97, Arg101, and Asn102 α4 and α5). A single point mutation in SlmA (R73D) was identified to be defective for FtsZ regulation *in vivo* and *in vitro* [20]. A later structural study of SlmA from *V. cholerae, E. coli*, and *K. pneumoniae* in complex with DNA and the conserved CTP of FtsZ confirmed this interaction interface. This study also located the binding pocket for FtsZ CTP on the SlmA surface that is only formed when the protein is bound to SBS [27], linking DNA binding of SlmA to its activity towards FtsZ.

The interaction between SlmA and the FtsZ CTP suggests that SlmA could compete with the membrane anchors ZipA and FtsA [26]. However, several observations indicate that the interaction with FtsZ’s CTP alone cannot explain SlmA’s inhibitory activity. First, SlmA inhibits FtsZ polymerization *in vitro* in the absence of membrane anchors [18]. Second, the cellular abundance of SlmA is far lower than that of FtsZ and its membrane anchors [18, 45, 46, 47], making such a competition model insufficient. Moreover, while sequestering proteins typically decrease the GTPase activity [37], SlmA does not change [48], and may even increase [17], FtsZ’s GTPase activity – behavior inconsistent with monomer sequestration. Together, these findings led to the proposal that SlmA acts through a secondary interaction that promotes filament destabilization at substoichiometric concentrations. Indeed, a weak interaction between SlmA and FtsZ’s globular domain has been described [16], but its structural basis as well as the functional consequences for FtsZ polymerization are not known.

To gain more insights into the mechanisms by which SlmA acts on FtsZ, we used an established *in vitro* assay based on purified proteins in fluorescent microscopy that allows directly watching treadmilling FtsZ filaments [49, 50, 8] and the effect of FtsZ-binding proteins [51, 52] in real time (**fig**. 1**b**). In the absence of SlmA, FtsZ and FtsA organize into a dynamic pattern of treadmilling filaments of rotating rings and moving streams on a surface of lipid bilayers [8, 53] to eventually reach a steady-state. Upon addition of WT SlmA in the presence of the SBS, we found that the fluorescence intensity corresponding to FtsZ dropped suddenly, before it slightly increased within 20 min to a new steady state (**fig**. 1**c,e**, **Supplementary Movie** 1). This effect was less pronounced in the absence of DNA, consistent with previous results [17]. When we repeated this experiment at different concentrations of SlmA, we found that this decrease in FtsZ intensity was concentration dependent (**fig**. S1**a,b**).

For comparability between individual experiments, we first quantified the decrease in fluorescence after addition of different SlmA concentrations (see **sec**. S3.3) and then fitted an inhibitory Hill function to the data to obtain a half-maximal effective concentration (EC_50_) for the different experimental conditions (**fig**. 1**g**. For WT SlmA in the presence of DNA, we found an EC_50_ of 79.3 ± 19.4 nM, about 15 times lower than the concentration of FtsZ and about 5 times lower than the concentration of FtsA used in our assay. In the absence of DNA this value increased to 427.2 ± 62.9 nM, consistent with the previous observation that DNA-binding promotes the formation of higher-order SlmA complexes with increased activity towards FtsZ [48, 27]. Next, we performed this experiment with SlmA R73D, previously described as inactive (**fig**. 1**d,f**, **Supplementary Movie** 2). We found that SlmA R73D still decreased the fluorescence intensity corresponding to FtsZ filaments on the membrane surface, although its EC_50_ increased to 511.2 ± 123.2 nM. In contrast to the wild-type protein, the activity of SlmA R73D did not depend on DNA (301.0 ± 165.5 nM)(**fig**. 1**f– i**,**fig**. S1**c,d**). Together, these experiments show that SlmA depolymerizes FtsZ filaments at concentrations well below that of FtsZ, and that binding to the SBS increases its activity about five-fold. Importantly, we also find SlmA R73D retains activity, with an EC_50_ about six times higher than that of the wild-type protein, supporting previous hypotheses for a secondary interaction with FtsZ next to its CTP.

To quantify filament turnover at different SlmA concentrations we performed fluorescence recovery after photo-bleaching experiments (FRAP). Assuming steady-state treadmilling with constant polymerization velocity and rapid monomer exchange, we hypothesize that the fluorescence recovery directly reports the mean residence time of subunits within filaments, where the fluorescence half-recovery time (*τ*_1/2_) represents a readout for the average filament length. In our experimental conditions with [FtsZ] = 1.25 µM and [FtsA] = 0.4 µM, we find *τ*_1/2_ to be 10.45±0.48 s (n=4) and the treadmilling velocity to be 53.40±7.19 nm s^*™*1^ (n=6), corresponding to a mean filament length of around 550 nm and consistent with what has been reported from HS-AFM experiments [53]. The presence of SlmA led to a marked decrease by 40% in FRAP recovery times even at SlmA concentrations lower than 100 nM, consistent with reduced subunit residence times and shortening of filaments (**fig**. 1**h**). At higher SlmA concentrations, however, recovery kinetics reached a plateau and became largely insensitive to further increases in SlmA, despite continued reductions in total polymer mass. At the same time, we found that the mean treadmilling velocity at these concentrations stayed constant (**fig**. 1**i**). This behavior suggests that SlmA initially affects filament length in a concentration-dependent manner, but at higher concentrations reaches a new regime in which a shorter steady-state filament length is established. In this regime, additional SlmA primarily reduces the number of filaments rather than further decreasing their length, thereby decoupling filament kinetics from total polymer abundance.

Together, this confirms the regulatory activity of SlmA, the ability of DNA to increase this activity and the role of SlmA R73. In addition, our observations support the catalytic activity of SlmA, with an EC_50_ 15 times lower than the FtsZ concentration. Finally, our data shows that mutating the CTP binding site on SlmA is not sufficient to completely abolish the activity, substantiating the previous hypothesis of a secondary interaction and two-pronged mechanism by which SlmA destroys FtsZ filaments.

### 2.2 SlmA behaves similar to FtsZ L272E, a bona fide capper

Previous SAXS experiments suggested an interaction of SlmA’s helices α4 and α7 with the C-terminal domain of FtsZ [16], which is distinct from FtsZ CTP-binding site with R73 on α4. Instead, this binding site corresponds to the bottom-surface of FtsZ and overlaps with the longitudinal interaction interface within the FtsZ filament. Binding of SlmA to FtsZ’s bottom-surface could therefore block the growing end of the filament inhibiting polymerization [21].

To benchmark the influence of a known capping protein, we compared SlmA-dependent filament regulation to that of a previously characterized filament end-binding mutant, FtsZ L272E, a variant of FtsZ reported to impair cell growth at substoichiometric concentrations by binding to filament growing ends and blocking further subunit addition [38] (**fig**. 2**a**). In our assay, addition of FtsZ L272E to filament patterns of WT FtsZ produced a strong and efficient reduction in filament intensity (**fig. 2b**, fig. S2) even at sub-equimolar concentrations, with titration experiments yielding an EC_50_ of 232.1 ± 116.0 nM, which lies between the EC_50_ values observed under different SlmA conditions (**fig**. 2**c,d**). Notably, FRAP analysis of FtsZ L272E revealed a biphasic response highly similar to that observed for SlmA, characterized by an initial decrease in recovery half-time at low concentrations followed by a plateau at higher concentrations despite continued reductions in total polymer mass (**fig**. 2**e**), while the mean treadmilling velocity stayed constant [50] (**fig**. 2**f**). This shared phenomenology suggests that SlmA and FtsZ L272E use the same underlying mechanism of filament end inhibition and support the interpretation that SlmA functionally acts as a filament capper under the conditions tested.

**Figure 2:**
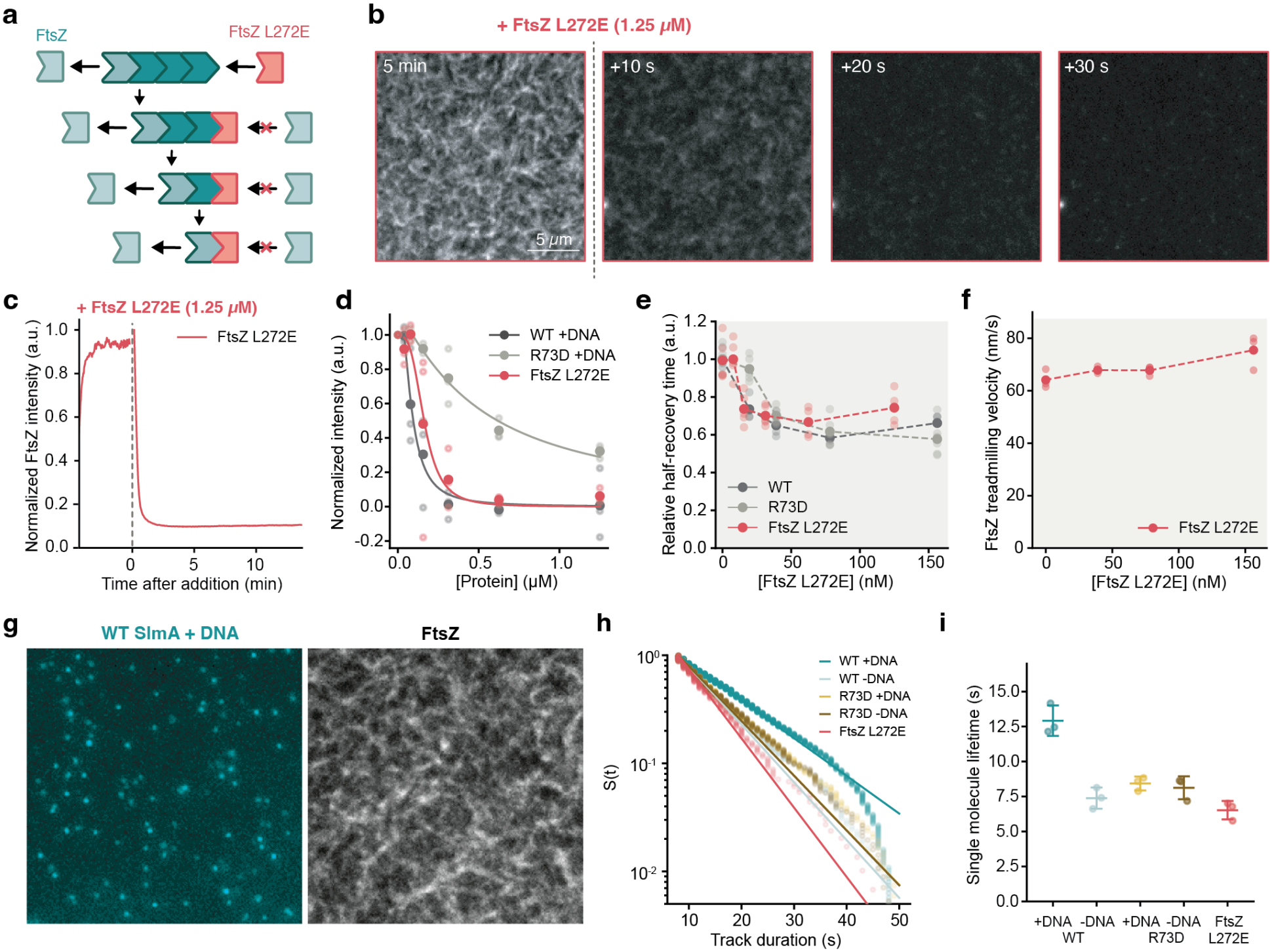
FtsZ L272E shows similar behavior as SlmA. **a)** The bottom-face mutant FtsZ L272E blocks polymerization efficiently destroying FtsZ filaments. **b)** Addition of 1.25 µM FtsZ L272E rapidly depolymerizes treadmilling filaments of FtsZ WT. **c)** Normalized fluorescence intensity corresponding to micrographs shown in b. The curve is representative of *n* = 3 independent experiments. **d)** Decrease of FtsZ fluorescence intensity (**sec. S3.3**) is FtsZ L272E concentration dependent. Curves correspond to fitted inhibitory Hill functions. **e)** Addition of FtsZ L272E decreases half-recovery time and increases filament turnover before reaching a plateau. At the same time, the treadmilling velocity **f)** stays constant. Results for WT SlmA and SlmA R73D are shown in black and dark grey, respectively. In d)–f), each dot represents one independent experiment (*n* = 3 per condition). **g)** Micrograph showing individual WT SlmA molecules labeled in Cy5 and filament pattern of Alexa488-labeled FtsZ filaments. **h)** Single molecule lifetime distribution and **i)** mean lifetimes for different versions of SlmA and FtsZ L272E. Curves in h) are representative of *n* = 3 independent experiments per condition. In i), dots represent individual experiments; bars show mean ± SD (*n* = 3).

To directly compare how different versions of SlmA and the bona-fide filament capping mutant FtsZ L272E interact with FtsZ filaments, we performed single-molecule imaging experiments. The total concentration of these inhibiting proteins we used in these experiments was in the nanomolar range, at which the FtsZ filament pattern remained unperturbed. In contrast to bulk biochemistry experiments for studying SlmA-FtsZ interaction, such as pelleting assays with stable filaments of severely hydrolysis-deficient FtsZ D212N [20], this single-molecule approach is sensitive enough to resolve even transient binding events, in particular under conditions where filaments undergo rapid treadmilling. This approach therefore provides a sensitive, direct readout of how long individual molecules remain associated with treadmilling filaments on the membrane surface, enabling quantitative comparison of protein-filament interactions under identical conditions.

First, and in contrast to previous reports [54], we found that under these conditions SlmA did not bind to the membrane on its own, as we detected single SlmA molecules only in the presence of FtsZ filaments (**fig**. S3, **Supplementary Movie** 3) showing that SlmA recruitment was FtsZ dependent. Using single-molecule tracking [49], we then quantified the residence times of individual Cy5-labeled wild-type SlmA and the R73D mutant in the presence and absence of DNA, as well as of FtsZ L272E in a background of FtsZ WT and FtsA (**fig**. 2**g-i**, **Supplementary Movie** 4). We find that WT SlmA in the presence of DNA exhibits a residence time of approximately 12 s, which exceeds the typical residence time of FtsZ WT monomers (**fig**. 2**h, i**) [8, 53]. In the absence of DNA, SlmA residence time decreases to ∼7.5 s, closely matching both FtsZ WT and the capping reference FtsZ L272E. Strikingly, SlmA R73D shows a similar residence time to WT SlmA without DNA and is not enhanced by DNA binding, indicating that the R73-mediated interaction alone cannot account for the observed binding to FtsZ filaments on the membrane.

Together, these measurements demonstrate that DNA binding strongly enhances the effective interaction lifetime of SlmA, consistent with DNA-dependent formation of higher-order oligomeric assemblies that increase avidity toward filament ends. The comparable residence times of SlmA R73D and FtsZ L272E further indicate that SlmA retains an additional interaction interface with filament subunits next to the previously proposed R73-dependent contact.

### 2.3 SlmA uses a second binding site to interact with the bottom surface of FtsZ

To identify the molecular basis of the additional SlmA-filament interaction inferred from our single-molecule measurements, we next sought to resolve the structural bases for SlmA’s activity. As a high-resolution structure of DNA-bound SlmA dimer with full-length FtsZ was not available, we used AlphaFold 3 [55] to predict the structure of this complex. The model, with an ipTM score of 0.84, robustly recapitulated the previously established interaction between the SlmA surface surrounding R73 and the conserved CTP of FtsZ (**fig**. 3**b,c**), consistent with prior biochemical and structural studies [20, 27]. In addition, AlphaFold 3 predicted a second, spatially distinct interface involving a region of SlmA centered around helix α7 that contacts the bottom face of FtsZ, which we termed the bottom interaction surface (BIS). Removing the FtsZ CTP from the prediction left the BIS unchanged and the model high-confidence (ipTM = 0.82), so the second interface does not require the canonical CTP contact. Interestingly, the bottom surface of FtsZ, which corresponds to the growing end of the filament, is also targeted by other known regulatory proteins such as MciZ [21], MipZ [33], and MinC [29], suggesting the growing end as a common target for FtsZ regulators (**fig**. S4).

**Figure 3:**
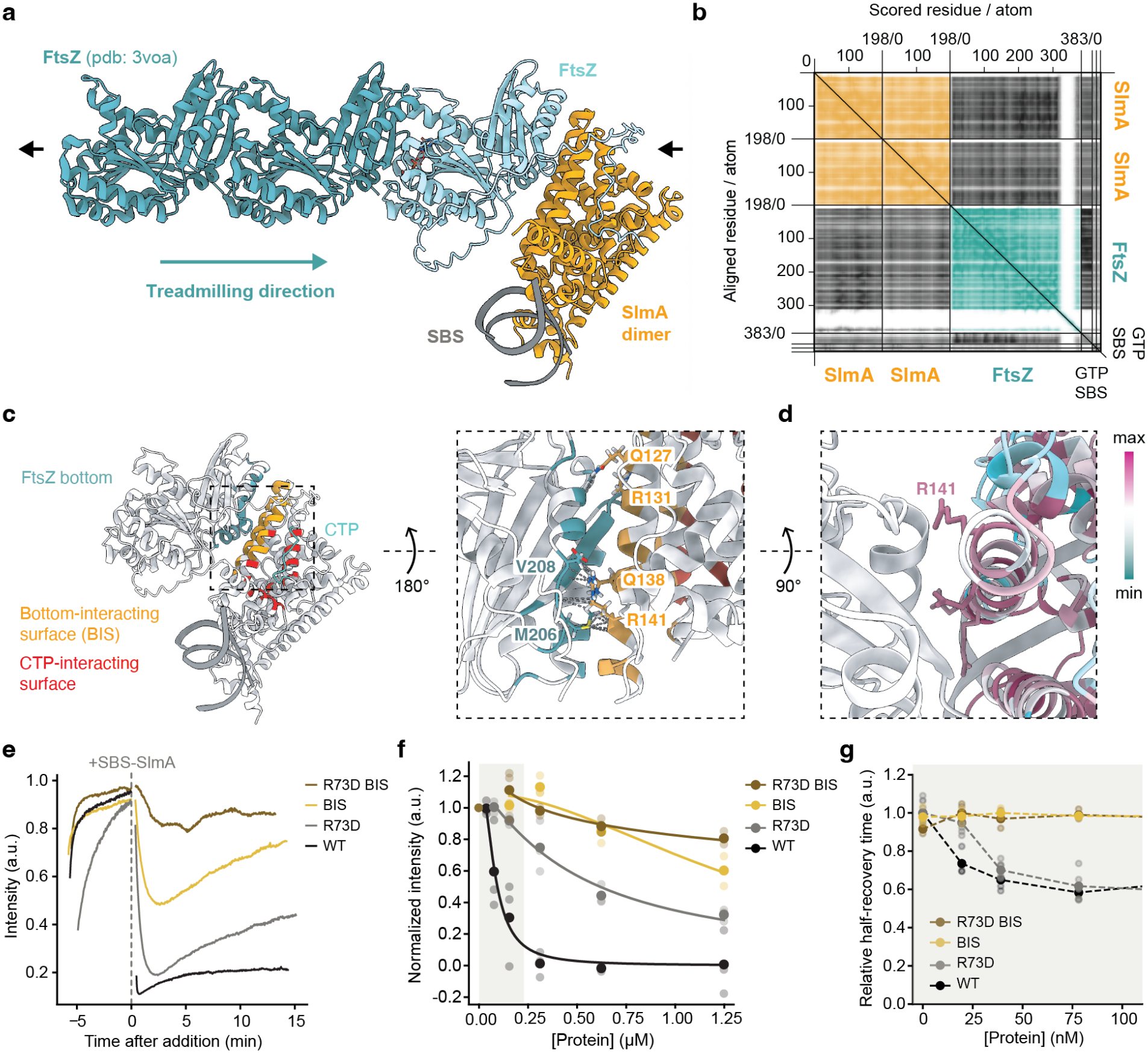
A new SlmA binding site for the interaction with FtsZ. **a)** AlphaFold 3 prediction of *Escherichia coli* FtsZ (light teal) matched onto the crystal structure of *Staphylococcus aureus* FtsZ in its filament conformation (dark teal, PDB 3VOA). **b)** Corresponding predicted aligned error (PAE) map. Dark grey areas correspond to predicted interaction between the SBS-SlmA dimer and FtsZ-GTP. **c)** Left: Predicted interaction clusters of the SBS-SlmA-FtsZ complex, teal, yellow = bottom interaction surface on FtsZ and SlmA, respectively; red = groove on SlmA interacting with FtsZ CTP. Within dashed box: SlmA R141 is predicted to interact with the T7 loop of FtsZ. **d)** Evolutionary conservation of amino acids as estimated by ConSurf, overlaid onto the predicted structure of SlmA. **e)** Normalized fluorescence intensity after addition of different versions of SlmA. Each trace was normalized to its projected maximum intensity. Traces are aligned to the time of SlmA addition (*t* = 0) and are representative of *n* = 3 independent experiments per condition. SBSSlmA BIS and SBS-SlmA BIS R73D show lower activity towards FtsZ. **f)** Decrease of FtsZ fluorescence intensity after SBS-SlmA addition. Curves correspond to fitted inhibitory Hill functions. **g)** Half-recovery times of FtsZ fluorescence after addition of SBS-SlmA BIS (yellow) and BIS R73D (brown). Curves for SBS-WT SlmA and R73D (black and dark grey) are shown as comparison. In f) and g), each dot represents one independent experiment (*n* = 3 per condition).

We further analyzed the predicted protein-protein complexes using two different computational tools, Pickluster [56], and LIVIA [57]. Both tools confidently resolved two distinct interaction clusters on SlmA. The first corresponds to the interface described in previous structural and biochemical studies [20, 27, 42], in which a hydrophobic groove on SlmA (13–105) engages the FtsZ CTP (371–379) (**fig**. 3**c** and **fig**. S4). The second cluster defines an interface in which SlmA residues on helices α4 and α7 contact the bottom surface of FtsZ, spanning FtsZ residues 205–219 and 287–296 through predominantly electrostatic interactions. FtsZ 205–219 contains the conserved T7 loop (GXXNXD, residues 204-209) that mediates longitudinal contacts between FtsZ subunits during polymerization [58]. SlmA contacts these T7 loop residues, in particular FtsZ M206 and V208 (**fig**. 3**c**), blocking interactions normally made by an incoming FtsZ subunit at the longitudinal polymerization interface. This interaction between SlmA and FtsZ is consistent with a filament end-capping mechanism. Consistent with a functional role for this surface, ConSurf [59] analysis showed that the SlmA residues forming the BIS are conserved, with Q127 and Q138 essentially invariant across SlmA orthologues and R131 and R141 conserved as basic residues (**fig**. 3**d**).

Next, to functionally validate the newly identified interaction surface of SlmA, we generated alanine substitution mutants targeting these conserved residues in the BIS of SlmA while leaving the CTP-binding groove intact. We first tested a SlmA double mutant (Q127A and R131A) (**fig**. S6), which retained wild-type activity, indicating that these two residues alone do not mediate functional interaction with the BIS. Similarly, a SlmA Q138A and R141A double mutant showed activity only slightly lower than the wildtype protein (**fig**. S6). We therefore generated a quadruple alanine substitution mutant (Q127A, R131A, Q138A, R141A = SlmA BIS), which resulted in a marked ∼15-fold increase in EC_50_ (1.26 ± 0.33 µM) (**fig**. 3**e,f**, **Supplementary Movie** 5). When we combined these mutations with R73D to disrupt both the canonical and novel interfaces simultaneously, we found a near complete loss of activity (EC_50_ = 3.26 ± 0.77 µM, **Supplementary Movie** 6). FRAP analysis revealed no detectable changes in turnover kinetics across these mutants (**fig**. 3**g**), and treadmilling velocity remained unchanged in all conditions tested. We also aimed to quantify the residence time of both, fluorescently labeled SlmA R73D-BIS and SlmA BIS on membrane-bound FtsZ filaments, but could not detect any binding events for these versions. These results indicate that the novel interaction interface on SlmA is essential for DNA-enhanced engagement of FtsZ subunits. The additive effect of combining BIS and R73D mutations is consistent with independent contributions of the two interfaces and suggest that the two interfaces likely act cooperatively to enable multivalent engagement of SBS-bound SlmA dimers with FtsZ filament ends.

### 2.4 BIS substitutions abolish the cell division regulation of SlmA

To determine whether BIS substitutions affect the function of SlmA to regulate *E. coli* cell division, we expressed SlmA BIS and SlmA R73D BIS in HC278, a *ΔslmA* strain in which minCDE expression is controlled by an arabinose-inducible promoter [20]. Functional complementation was examined under Min^-^ SlmA^-^ conditions by inducing SlmA variant expression with 0.1 mM IPTG. We confirmed that expression of WT SlmA fully restored colony formation on LB agar and restored the WT phenotype of the HC278 cell in absence of arabinose, indicating that this level of induction was sufficient to complement the loss of endogenous SlmA as found previously. In contrast, neither SlmA R73D, SlmA BIS nor SlmA R73D BIS restored plating efficiency or normal cell morphology (**fig**. 4**a**). Instead, all three variants exhibited severe plating defects and extensive filamentation similar to those observed with the empty-vector control and the previously characterized inactive mutant SlmA R73D [18], indicating a complete loss of SlmA function *in vivo*.

**Figure 4:**
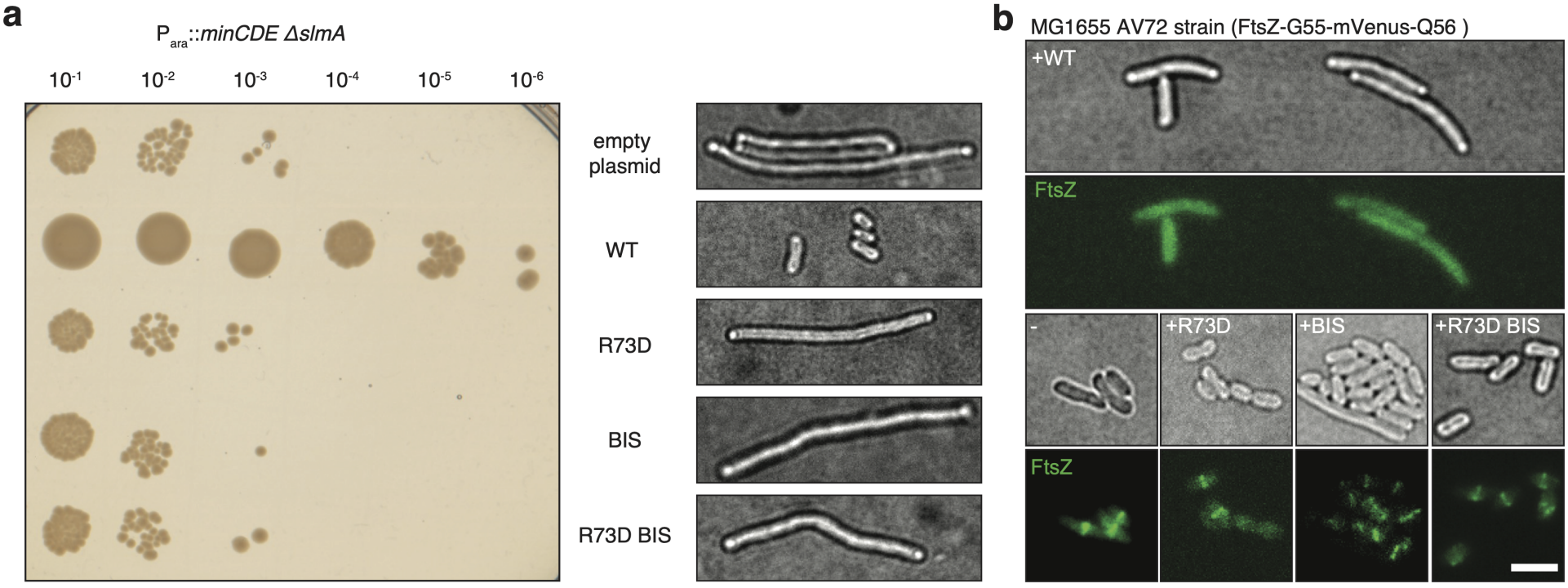
The bottom-interaction interface of SlmA plays an important role for Z-ring regulation a) Left panel: Overnight cultures of HC278 (*Para::minCDE ΔslmA*) carrying complementation plasmids expressing the indicated SlmA variants were normalized to the same OD_600_ of 0.2, serially diluted, and 5 µL of each dilution was spotted onto LB agar plates supplemented with 0.1 mM IPTG to induce expression of the SlmA variants. **Right panel:** To examine cell morphology, overnight cultures of the same strains were reinoculated into LB broth and grown to an OD_600_ of 0.5–0.6. Cultures were then induced with 0.1 mM IPTG for 2 h. Samples were collected and examined using DIC optics. Representative images are shown. **b)** Overnight cultures of MG1655 AV72 (*FtsZ-G55-mVenus-Q56*) carrying complementation plasmids were reinoculated into LB broth and grown to an OD_600_ of 0.5–0.6. Cultures were induced with 1 mM IPTG for 2 h, and samples were imaged using DIC and green fluorescence microscopy. Representative images are shown. Scale bar, 5 µm. All experiments were performed independently at least three times (*n* ≥ 3).

To further assess the division inhibitory activity of these variants, we examined their effects on FtsZ-ring assembly using the MG1655 AV72 strain, in which FtsZ-G55-mVenus-Q56 is the sole chromosomal copy of FtsZ. Consistent with previous reports, induction of wild-type SlmA with 1 mM IPTG completely disrupted Z-ring assembly, resulting in the absence of detectable midcell FtsZ rings [18]. In contrast, expression of either SlmA BIS or SlmA R73D BIS failed to perturb Z-ring formation, and cells retained normal midcell FtsZ rings similar to those previously reported for the inactive SlmA R73D mutant (**fig**. 4**b**).

Collectively, these results demonstrate that both the BIS and R73D BIS substitutions abolish the division regulatory activity of SlmA, identifying the corresponding residues as essential determinants of SlmA-mediated inhibition of bacterial cytokinesis.

### 2.5 Simulations demonstrate effective filament regulation via end-capping

Our experiments indicate that SlmA caps the growing ends of treadmilling FtsZ filaments, thereby suppressing further elongation. Other FtsZ regulators appear to act through similar end-capping mechanisms [21, 33, 29], raising the question about the advantage of this strategy compared to alternative modes of regulation. To address this, we developed a stochastic model of FtsZ treadmilling and compared growing-end capping with monomer sequestration and filament severing [60] in terms of their effect on polymer mass and the regulator concentration required.

We simulated treadmilling FtsZ filaments coupled to a finite, well-mixed pool of monomers and regulators using the Gillespie algorithm (**Supplementary Movie** 7). In this simplified representation, treadmilling filament dynamics are fully described by four parameters: the rate constant of monomer attachment at the growing end (*k*_on_), the rate constant of GTP hydrolysis within the filament (*k*_hyd_), the rate constant of monomer detachment at the shrinking end (*k*_off_), and the nucleation equilibrium constant (*K*_nuc_) (**fig**. 5**a I**) [37]. The full model and its assumptions are described in Section 4.8.

**Figure 5:**
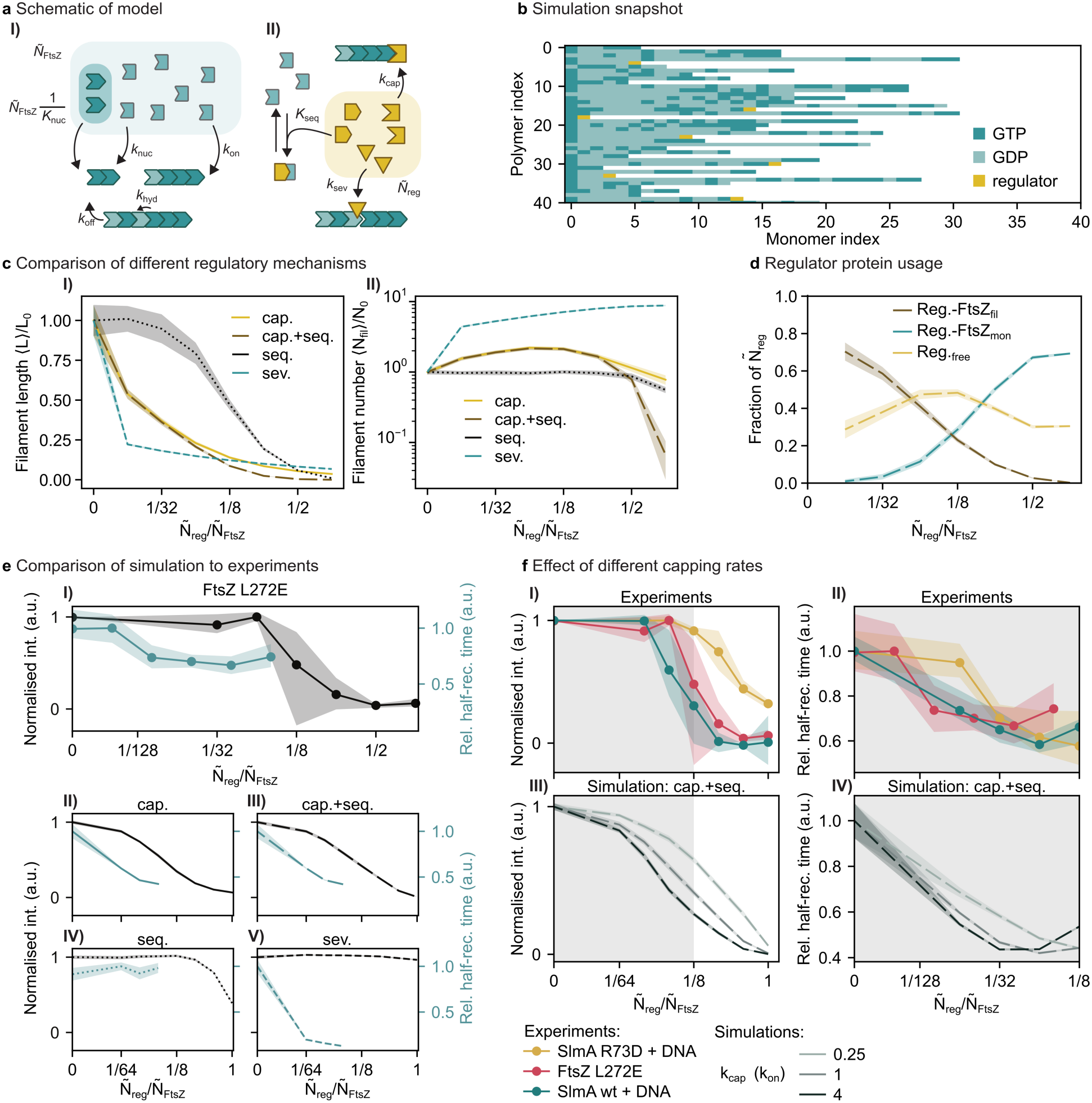
Simulations identify growing-end capping as an efficient mechanism for regulating treadmilling FtsZ filaments. **a)** Schematic of the model. **I** Filaments nucleate from nucleation-competent free monomers (fraction 1/*K*_nuc_), grow at the growing end (*k*_on_), hydrolyse GTP (*k*_hyd_) and lose GDP-bound subunits at the shrinking end (*k*_off_). **II** Regulators cap growing ends (*k*_cap_), sequester free monomers (*K*_seq_) or sever filaments (*k*_sev_). **b)** Simulation snapshot with capping regulators at *Ñ*_reg_/*Ñ*_FtsZ_ = 1/8 (**Supplementary Movie** 7). Each row is one filament, drawn from the shrinking end (monomer index 0) to the growing end; subunits are coloured by nucleotide state, regulators in yellow. **c)** Mean filament length ⟨*L*⟩ (**I**) and mean filament number ⟨*N*_fil_⟩ (**II**), relative to their unregulated values *L*_0_ and *N*_0_, as a function of the regulator ratio *Ñ*_reg_/*Ñ*_FtsZ_ for capping (cap.), combined capping and sequestration (cap.+seq.), sequestration (seq.) and severing (sev.). **d)** Fate of regulators in the combined mechanism: fraction of *Ñ*_reg_ capping filament ends (Reg.-FtsZ_fil_), sequestering free monomers (Reg.-FtsZ_mon_) or free (Reg._free_). **e)** Normalised fluorescence intensity (black, left axes) and relative half-recovery time (teal, right axes), measured for FtsZ L272E (**I**) and simulated with *k*_cap_ = *k*_on_ for capping (**II**), capping with sequestration (**III**), sequestration (**IV**) and severing (**V**). **f)** Effect of the capping rate constant on the same two quantities (**I, III**: intensity; **II, IV**: half-recovery time), measured for WT SlmA + DNA, SlmA R73D +1D3 NA and FtsZ L272E (**I, II**) and simulated for capping with sequestration at *k*_cap_/*k*_on_ = 0.25, 1 and 4 (**III, IV**). Grey shading in I and III marks the range shown in II and IV.

We first fitted the unregulated model to experimentally determined monomer lifetime distributions [8, 51], treadmilling speeds, and FRAP half-recovery times [51, 53, 52] using Bayesian optimisation [61] (**fig**. S9**a**–**c**; Table S4). A parameter scan confirmed the robustness of this fit (**fig**. S10) and showed that reproducing the observed treadmilling behaviour requires *k*_hyd_ to be between 1 s^*™*1^ to 100 s^*™*1^, orders of magnitude higher than assumed in previous models of treadmilling filaments [37, 62].

We then implemented four regulatory mechanisms (**fig**. 5**a II**): pure capping, in which regulators bind and block the growing end with the rate constant *k*_cap_ (**fig**. 5**b**); pure sequestration, in which regulators bind free monomers, controlled by the sequestration equilibrium constant *K*_seq_; combined capping and sequestration, as end-binding proteins could also interact with FtsZ monomers in solution; and filament severing, a mechanism previously suggested for MinC [60], with the per-monomer-monomer-bond rate constant *k*_sev_. For each mechanism, we simulated a finite pool of *Ñ*_FtsZ_ monomers with increasing numbers of regulators, *Ñ*_reg_, expressed as the regulator ratio *Ñ*_reg_/*Ñ*_FtsZ_.

We first assumed that a free regulator caps a growing end with the same rate constant per molecule as a free monomer attaches to it, *k*_cap_ = *k*_on_*Ñ*_reg_/*Ñ*_FtsZ_ (sec. 4.8.7), and that sequestration and severing happen on similar timescales (sec. 4.8.8 and 4.8.9). Under this assumption, all four mechanisms reduced the mean filament length, ⟨*L*⟩, relative to the unregulated condition, *L*_0_, but sequestration alone required an approximately tenfold higher regulator concentration to achieve a comparable reduction (**fig**. 5**c I**). The mechanisms differed markedly in their effect on the mean number of filaments, ⟨*N*_fil_⟩ (**fig**. 5**c II**). Capping, alone or combined with sequestration, slightly increased the number of filaments at substoichiometric regulator concentrations, but reduced it once regulator concentrations approached that of FtsZ. Sequestration alone had little effect on filament number, whereas severing caused a marked increase. Thus, at substoichiometric concentrations, only capping and severing substantially reduced filament length, and only capping did so while keeping filament number close to that of the unregulated system (**fig**. 5**c**).

Capping was substantially more effective than monomer sequestration. At a given regulator ratio, capping alone reduced filament length far more than sequestration alone, and adding sequestration to capping changed both quantities only slightly (**fig**. 5**c**). In the combined mechanism, regulators preferentially capped existing filaments at low regulator ratios. At ratios 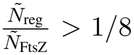, the number of free regulators began to decrease as regulators bound the free FtsZ monomers (**fig**. 5**d**), consistent with our observation of two activity regimes in our FRAP experiments (**fig**. 1**h**, **fig**. 2**e**).

Simulations with capping, alone or combined with sequestration, qualitatively reproduced the decreases in fluorescence recovery time and signal intensity observed for the bona fide capper FtsZ L272E (**fig**. 5**e I, II** and **III**). Sequestration alone showed no decrease in half-recovery time and only decreased fluorescence intensities at high regulator fractions 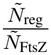 in our simulations (**fig**. 5**e IV**). Severing decreased half-recovery time at low regulator fractions 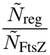 but conserves fluorescence intensity even at high regulator fractions (**fig**. 5**e V**).

The assumption *k*_cap_ = *k*_on_ is best justified for FtsZ L272E, because this mutation affects only the growingend interface involved in monomer addition. Even so, we do not expect this simplified model to reproduce the measured recovery times quantitatively, because neither *k*_cap_ nor the lifetimes of regulator-monomer complexes could be determined experimentally (see Sections 4.8.7 and 4.8.8 and SI Section S3.2). Indeed, varying *k*_cap_ relative to this reference value improved the agreement between simulated and measured fluorescence intensities (**fig**. 5**f I** and **III**) and recovery times (**fig**. 5**f II** and **IV**). At higher regulator concentrations, the combined mechanism reproduced the slight increase in recovery time observed experimentally (**figs**. 2**e** and 5**f IV**). The distinct behaviours of FtsZ L272E, WT SlmA + DNA, and the less active SlmA R73D + DNA could all be reproduced by varying only the capping activity. A more quantitative agreement could likely be obtained by fitting *k*_cap_, and potentially *K*_seq_, directly to the experimental data.

Together, these simulations identify growing-end capping as an efficient mechanism for regulating treadmilling filaments. Capping recapitulates the defining features of SlmA activity: a strong reduction in filament length at substoichiometric concentrations, accompanied by only a slight increase in filament number. Severing also shortened filaments but markedly increased their number, which is inconsistent with our experimental observations. Structural predictions of the SlmA-FtsZ complex further argue against severing, because SlmA binds the growing-end interface of FtsZ and cannot engage subunits within the filament (**fig**. 3). The model suggests that capping dominates at low regulator concentrations, whereas sequestration likely contributes at high concentrations. Moreover, differences between FtsZ L272E, WT SlmA, and SlmA R73D reflect primarily differences in effective capping activity. Because a single capping protein can inactivate an entire filament, end-capping allows substoichiometric amounts of regulator to control FtsZ polymerization while preserving treadmilling dynamics, which may explain why this strategy has been adopted by multiple FtsZ regulators.

## 3 Discussion

Negative regulators of FtsZ from different species have been identified and were proposed to work with different mechanisms [63, 64, 65, 41, 26]. New evidence showed that a common theme for the regulation of FtsZ polymerization emerges: all these negative regulators target the growing plus end of the FtsZ filament, the polymerization-competent, bottom surface of its C-terminal globular domain. Structural and biophysical studies, such as crystallography (MciZ [21]), NMR (MinC [29] ) and HDX-MS (MipZ, [33]), have helped to define how these factors engage with FtsZ providing evidence for a capping mechanism to control FtsZ polymerization. Despite this similarity in protein-protein interaction, the consequences of plus-end capping for the dynamics of a treadmilling filament, and how it translates into changes in filament mass has not been characterized. Here, we address this gap by studying how SlmA acts on FtsZ polymers under non-equilibrium, treadmilling conditions. Using *in vitro* reconstitution and direct visualization of treadmilling filaments by fluorescence microscopy, we show that SlmA acts as a catalytic inhibitor of FtsZ assembly. We find that SlmA operates at substoichiometric concentrations, shortens filaments and shows binding kinetics consistent with transient association of filament ends. Together with prior observations that SlmA accelerates filament disassembly without reducing FtsZ’s GTPase activity, these results are most consistent with a capping mechanism.

SlmA was previously found to interact with FtsZ by binding to FtsZ’s C-terminal peptide [26, 27], while several studies pointed out additional regulatory interactions are likely [16, 27]. Using protein structure predictions, we identify a previously unrecognized interface between SlmA and the bottom interface of FtsZ. Mutations designed to disrupt the newly identified interface strongly impair activity *in vitro* and abolish it *in vivo*, demonstrating that it constitutes the primary functional interface. In contrast, the R73D mutation in SlmA, which disrupts interaction with the FtsZ C-terminal peptide, only partially reduces activity *in vitro* suggesting that the C-terminal peptide contact modulates SlmA’s activity and is not essential. Single-molecule imaging revealed that WT SlmA had the longest lifetimes, whereas R73D showed reduced lifetimes and binding of the BIS mutants was undetectable, a pattern that closely correlates with the activities measured in our reconstitution assay.

Importantly, we find that the experimentally observed influence of SlmA on FtsZ filaments can be recapitulated in a minimal theoretical model. Simulations incorporating end-specific inhibition reproduce the key experimental observations, including the switch from steady-state treadmilling into net depolymerization and filament shortening at substoichiometric regulator concentrations. This agreement between experiment and theory supports a capping based mechanism and demonstrates that the observed effects emerge directly from perturbation of filament-end kinetics without invoking additional regulatory complexity.

These results clarify why plus-end targeting is such a productive regulatory strategy for treadmilling filaments. FtsZ treadmilling is a non-equilibrium steady state driven by GTP hydrolysis, in which subunit addition at the plus end and dissociation from the minus end proceed simultaneously. Because minus-end dissociation continues unimpeded, blocking plus-end growth therefore redirects an existing steady-state turnover into net shrinkage. A single capper therefore exerts disproportionate influence over filament mass as it exploits the kinetic polarity of treadmilling filaments. This stands in contrast to the regulation of isodesmic polymers, where the absence of end asymmetry means that a regulator targeting one face of a subunit has no privileged effect on filament dynamics. Instead, the regulator must engage a large fraction of subunits, or induce cooperative conformational changes, to achieve comparable control. The evolutionary convergence on plus-end targeting among structurally unrelated FtsZ regulators – SulA, MciZ, MinC, MipZ, and now SlmA – is, in this light, not coincidental. Treadmilling therefore makes filaments intrinsically more controllable than equilibrium polymers.

As a nucleoid-associated protein, SlmA locally suppresses Z-ring formation over the chromosome while preserving the availability of FtsZ for assembly in regions permissive for division. By capping growing filaments, SlmA can reduce FtsZ polymerization specifically where it is bound, thereby redistributing FtsZ away from these regions rather than globally depleting the monomer pool. In contrast, a sequestration-based mechanism would act independently of filament location and would therefore be less spatially specific and less efficient, as it would reduce the availability of FtsZ monomers throughout the cell.

Taken together, our findings establish a two-pronged mechanism for the inhibition of FtsZ. First, SlmA binds to FtsZ CTP via a groove on helices α4 and α5, second, it acts as a plus-end capper of FtsZ filaments by binding to the T7 loop via the opposite face of helix α4 and helix α7. As a result, SlmA can efficiently depolymerize treadmilling filaments at concentrations much lower than that of FtsZ. More broadly, our results establish a systems-level understanding of how capping operates on a treadmilling filament in ways that prior structural studies did not address. This principle of using kinetic polarity as a property to allow for controlling filament mass by blocking growth also plays a critical role for the alignment of FtsZ filaments during Z-ring maturation [62] and is likely to be relevant beyond SlmA and FtsZ in *E. coli*. Given that FtsZ-based division is also used by many archaea [66], it will be interesting to determine whether analogous mechanisms of filament regulation contribute to Z-ring formation in these species.

## 4 Material and methods

Bacterial strains, plasmids and growth conditions. All bacterial strains and plasmids are listed in **Table** 1. *E. coli* strains were grown at 37 ^*°*^C in Lysogeny broth (LB) medium with shaking (250 rpm) or on LB agar plates supplemented with the appropriate antibiotics (100 µg mL^*™*1^ ampicillin, 50 µg mL^*™*1^ kanamycin). *E. coli* DH5α was used for plasmids propagation. *E. coli BL21* (DE3) was used for overexpressing proteins. MG1655 AV72 strain (carrying chromosomally encoded FtsZ-G55-mVenus-Q56 as a sole FtsZ copy) was used for *in vivo* imaging of FtsZ ring [52]. *E. coli* HC278 [*Para::minCDEΔslmA*] was used to analyze the *in vivo* function of SlmA complementation in absence of arabinose [20].

**Table 1:** Strains and plasmids used in this study.

| Strains/Plas-mids | Genotype | Source/Reference |
|---|---|---|
| **Strains** *Escherichia coli* |  |  |
| DH5α | *F<sup>−</sup> hsdR17 deoR recA1 endA1 phoA supE44 thi-1 gyrA96 relA1 Δ(lacZYA-argF)U169 ϕ80dlacZΔM15* | Gibco BRL |
| BL21 (DE3) | *fhuA2 [lon] ompT gal (λ DE3) [dcm] ΔhsdS; λ DE3 = λ sBamHIo ΔEcoRI-B int::(lacI::P<sub>lacUV5</sub>::T7 gene1) i21 Δnin5* | New England Biolabs |
| HC278 | *rph1 ilvG rfb-50 ΔlacIZYA::frt (frt araC P<sub>ara</sub>)::minCDE ΔslmA::frt* | [67] |
| MG1655 AV72 | *rph-1 ilvG rfb-50 ftsZ::ftsZ-G55-mVenus-Q56* | [68] |
| **Plasmids** |  |  |
| pTB146 | Amp<sup>r</sup>, expression plasmid containing replicon ColE1 (pBR322) | [6] |
| pXY38 | *slmA* cloned into pTB146 by Gibson assembly | This study |
| pXY39 | *slmA<sup>R73D</sup>*, generated by site-directed mutagenesis using pXY38 as template | This study |
| pXY40 | *slmA<sup>BIS</sup>*, generated by site-directed mutagenesis using pXY38 as template | This study |
| pXY41 | *slmA<sup>R73D,BIS</sup>*, generated by site-directed mutagenesis using pXY38 as template | This study |
| pMAL-c-*SulA* | SulA expression plasmid under the control of a *tac* promoter | [69] |
| pBAD33-Kan | Contains a p15A origin of replication; low-to-medium copy number; suitable for complementation in *E. coli* | From our lab |
| pXY43 | Arabinose promoter of pBAD33-Kan replaced with the *lacI* gene and T7 promoter region from pTB146 by Gibson as-sembly | This study |
| pXY44 | *slmA* cloned into pXY43 by Gibson assembly | This study |
| pXY45 | T7 promoter of pXY44 replaced with the *tac* promoter region from pMAL-c-*SulA* by Gibson assembly | This study |
| pXY46 | *slmA<sup>R73D</sup>*, generated in pXY45 by site-directed mutagenesis for complementation of Δ*minCDE*Δ*slmA* | This study |
| pXY47 | *slmA<sup>BIS</sup>*, generated in pXY45 by site-directed mutagenesis for complementation of Δ*minCDE*Δ*slmA* | This study |
| pXY48 | *slmA<sup>R73D,BIS</sup>*, generated in pXY45 by site-directed mutagen-esis for complementation of Δ*minCDE*Δ*slmA* | This study |

### 4.1 Cloning

For SlmA overexpression, the wild-type (WT) *slmA* gene was cloned into the SUMO-tag-containing expression vector pTB146 [17, 8] using Gibson assembly [70]. The resulting plasmid encoded an N-terminally His_6_-SUMO-tagged SlmA fusion protein and was transformed into *E. coli* BL21 cells for protein expression. Point mutations were introduced into *slmA* by site-directed mutagenesis using the Q5 Site-Directed Mutagenesis Kit (NEB, USA).

To generate an IPTG-inducible, His_6_-tagged SlmA expression plasmid for *in vivo* functional characterization in *E. coli*, we constructed a derivative of pBAD33-Kan by sequential Gibson assembly. First, the native arabinose promoter of pBAD33-Kan was replaced with the *lacI* gene and the T7 promoter region, both amplified from pTB146, to generate pXY43. Next, the *slmA* coding sequence was PCR-amplified and inserted downstream of the T7 promoter in pXY43, yielding pXY44. Finally, the T7 promoter of pXY44 was replaced with the *tac* promoter amplified from pMAL-c-SulA, concurrently introducing an N-terminal His-tag in frame with SlmA, to yield pXY45. All constructs were verified by Sanger sequencing to confirm sequence integrity, correct promoter replacement, and reading frame continuity.

### 4.2 Protein purification

Wild-type SlmA and mutant proteins were expressed in *E. coli* BL21 cells grown in LB medium supplemented with ampicillin (100 µg mL^*™*1^; Sigma-Aldrich) at 37 ^*°*^C to an OD_600_ of 0.5–0.6. Protein expression was induced with 1 mM IPTG, followed by overnight incubation at 20 ^*°*^C with shaking at 220 rpm. Cells were harvested by centrifugation at 8000 ×g for 20 min at 4 ^*°*^C and resuspended in lysis buffer (50 mM Tris-HCl, pH 8.0; 150 mM NaCl) containing protease inhibitors (Roche) and DNase I (Sigma-Aldrich). Cells were lysed by sonication on ice, and the lysate was clarified by centrifugation at 20 000 ×g for 30 min at 4 ^*°*^C. The supernatant was applied to a Ni-NTA gravity-flow column (Thermo Scientific) equilibrated with lysis buffer. After washing with lysis buffer containing 20 mM imidazole, bound proteins were eluted with lysis buffer containing 300 mM imidazole. To remove the SUMO tag, eluted proteins were incubated with SUMO protease and dialyzed overnight at 4 ^*°*^C against lysis buffer. The cleavage mixture was then passed over a second Ni-NTA column (Thermo Scientific), and the cleaved protein was recovered in the flow-through. Residual imidazole was removed on a PD-10 desalting column (GE Healthcare), and proteins were eluted in storage buffer (50 mM Tris-HCl, pH 8.0; 150 mM NaCl; 10 % glycerol). Protein purity was assessed by SDS-PAGE. Purified proteins were aliquoted and stored at ™70 ^*°*^C until use. FtsA and FtsZ (wild-type and L272E mutant) were expressed and purified as previously described [8].

### 4.3 Fluorescence microscopy

TIRF imaging was performed on a Visitron iLAS2 TIRF microscope equipped with a 63× Olympus oilimmersion objective (NA 1.46) and a quad-band laser filter set (405 nm, 488 nm, 561 nm and 640 nm). Fluorescence signals were detected with a Photometrics Evolve 512 EMCCD camera (512 × 512 pixels, 16 µm pixel size), giving an effective pixel size of 0.1 µm in the sample plane.

Time-lapse images were acquired with an EM gain of 200, an exposure time of 50 ms, a frame interval of 2 s and a laser power of 4 % to 10 %. For fluorescence recovery after photobleaching (FRAP) experiments, images were acquired with an EM gain of 200 at a frame interval of 1 s, comprising 5 pre-bleach and 195 post-bleach frames. Photobleaching was performed at a laser power of 40 % to 60 %. Single-molecule images were acquired with an EM gain of 200, an exposure time of 50 ms, a frame interval of 2 s and a laser power of 60 % to 80 %.

### 4.4 Image analysis

Treadmilling speed, FRAP and single-particle tracking were analyzed from TIRF movies as described previously, using ImageJ plug-ins and custom Python scripts [51, 52, 53].

### 4.5 Spotting assays

*E. coli* strain HC278 carrying pXY43-derived plasmids encoding the indicated SlmA variants was grown in LB medium supplemented with 50 µg mL^*™*1^ kanamycin at 37 ^*°*^C. Overnight cultures were normalized to an OD_600_ of 0.2 and serially diluted tenfold. Five microliters of each dilution were spotted onto LB agar plates supplemented with kanamycin (50 µg mL^*™*1^) and IPTG (0.1 mM) to induce expression of the SlmA variants. Plates were incubated overnight at 30 ^*°*^C and imaged the following day.

### 4.6 Agar pad preparation for *in vivo* imaging

To assess the effect of SlmA variant expression on cell shape (*E. coli* strain HC278) or on the Z-ring (strain MG1655 AV72), overnight cultures were diluted 1:100 (v/v) into fresh LB medium and grown at 37 ^*°*^C to mid-log phase (OD_600_ of 0.5–0.6). Protein expression was induced with IPTG, and cultures were incubated for a further 2 h at 30 ^*°*^C. Cells were then diluted 1:100 in LB, immobilized on 1.2 % agarose pads and imaged by bright-field and fluorescence microscopy.

Induction levels differed between the two assays: complementation of the Min^−^ SlmA^−^ defect was tested under low-level induction (100 µM IPTG), whereas FtsZ ring disruption in wild-type *E. coli* was assessed under high-level induction (1 mM IPTG).

### 4.7 Structural prediction and interface analysis

#### 4.7.1 Structural prediction

The structure of the *E. coli* SlmA dimer in complex with FtsZ, GTP and its cognate double-stranded SlmA binding site (SBS) DNA was predicted with AlphaFold 3 [55] using the AlphaFold Server (https://alphafoldserver.com). The input comprised the full-length sequences of two SlmA chains (UniProt P0C093) and one FtsZ chain (UniProt P0A9A6), GTP as a ligand, and the double-stranded DNA sequence of an SBS from the *E. coli* chromosome. The top-ranked model, as judged by the predicted interface TM-score (ipTM *>* 0.85), was selected for further analysis.

#### 4.7.2 Clash removal

Prior to interface analysis, hydrogen atoms and partial charges were added to the predicted model with the addh and addcharge commands in UCSF ChimeraX [71]. The DNA and GTP components were excluded to avoid parameterization issues, and the protein chains were geometry-minimized with the ISOLDE plugin [72] using the AMBER ff14SB force field until steric clashes were resolved.

#### 4.7.3 Interface analysis

Residue–residue contacts at the SlmA–FtsZ interface were identified with the Pickluster tool [56] in UCSF ChimeraX. Contacts were evaluated from van der Waals overlap values, where positive values indicate steric clashes and negative values favorable van der Waals contacts. The interface residues were then partitioned into two interaction clusters according to their spatial distribution: one engaging the FtsZ C-terminal region (Tyr371–Arg379) with a hydrophobic patch on SlmA, and a second engaging FtsZ residues 205–296 with a predominantly electrostatic surface on SlmA. Contacts between specific residue subsets were analyzed further with the contacts command in UCSF ChimeraX using default van der Waals overlap parameters.

Interface quality was assessed with LIVIA [57], which derives a set of confidence metrics for each chain pair based on the AFM-LIS framework [73]. For every inter-chain residue pair with a PAE of ≤12 Å, LIVIA converts the PAE into a local interaction probability; averaging these probabilities over all qualifying pairs yields the Local Interaction Score (LIS). Restricting the average to pairs that are also in physical contact (C*β*–C*β* distance ≤ 8 Å) gives the contact LIS (cLIS), and combining LIS and cLIS into a single value produces the integrated LIS (iLIS). Residues contributing to these interactions are classified accordingly as Local Interaction Residues (LIR, PAE ≤ 12 Å) or, when the stricter contact criterion is also met, as contactfiltered LIR (cLIR). LIVIA additionally calculates ipSAE and actifpTM scores directly from the PAE matrix and inter-chain contact data, alongside the platform-reported ipTM score, allowing comparison across several confidence metrics. For complexes with more than two chains, LIVIA computes all pairwise combinations of these metrics and presents them in a single summary score matrix.

### 4.8 Simulations

#### 4.8.1 Notation

Our Gillespie solver tracks integer numbers of molecules. *Ñ*_FtsZ_ is the total number of FtsZ molecules and *N*_FtsZ_ the number of FtsZ molecules that are not free monomers, i.e. polymerised in filaments or sequestered by a regulator. The number of free FtsZ monomers is therefore *Ñ*_FtsZ_ ™ *N*_FtsZ_. Analogously, *Ñ*_reg_ is the total number of regulators, *N*_reg_ the number of regulators bound to FtsZ, and *Ñ*_reg_ ™ *N*_reg_ the number of free regulators. A tilde always denotes a total.

Each process *i* occurs with a rate *r*_*i*_, the expected number of events of that process per second in the whole system. Each rate is calculated from a rate constant *k*_*i*_ in s^*™*1^ and the current state of the system. Rate constants of processes that consume free FtsZ monomers are defined for the case that all FtsZ is monomeric, *N*_FtsZ_ = 0; the rate is then scaled by the fraction of free monomers (*Ñ*_FtsZ_ ™ *N*_FtsZ_)/*Ñ*_FtsZ_. For example, a single uncapped filament end grows with rate *k*_on_(*Ñ*_FtsZ_ ™ *N*_FtsZ_)/*Ñ*_FtsZ_. The rate constants are the input parameters of the solver and are fitted to experimentally observed behaviour (**sec**. S3.4).

All rates depend on time *t* and the system state **x**, *r*_*i*_ = *r*_*i*_(*t*, **x**), which we omit for ease of notation. For any state variable *X*, ⟨*X*⟩ denotes its time average at steady state,

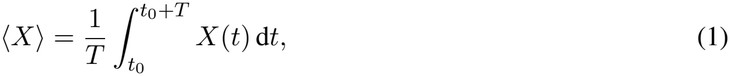

where *t*_0_ is a time after which the system has reached steady state and *T* is the averaging window.

Table S1 lists the input parameters of the model, Table S3 the state variables, and Table S2 the rates of all processes, which we derive in **sec**. 4.8.3–4.8.9.

#### 4.8.2 Rate calculation

Gillespie solvers have been used successfully to simulate systems of treadmilling filaments, both for single filaments [74, 75] and for multiple filaments [37]. For *n* possible processes with rates *r*_*i*_, *i* = 1, …, *n*, and total rate 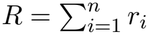, the solver draws the next timestep as

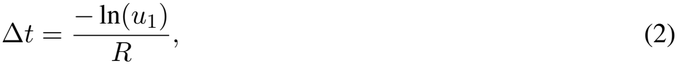

where *u*_1_ is a random number drawn uniformly from (0, 1]. Δ*t* is therefore exponentially distributed with mean 1/*R*. The solver then executes process *i* with probability *r*_*i*_/*R*. Our implementation considers all processes shown in **fig**. 5**a**.

#### 4.8.3 Nucleation

Nucleation of a new filament is a second-order process [37]: a free monomer in a nucleation-competent conformation binds a second free monomer. We assume that this conformation is in fast equilibrium with the inactive conformation, so that the number of nucleation-competent monomers is

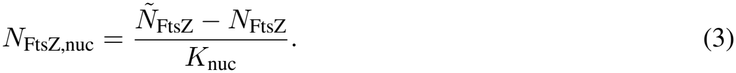

Eq. (3) defines the dimensionless nucleation equilibrium constant *K*_nuc_ ≥ 1 as the ratio of free monomers to nucleation-competent monomers, so that 1/*K*_nuc_ is the fraction of free monomers that are nucleation-competent. Each nucleation-competent monomer binds a free monomer with the same rate as an uncapped filament end (eq. (5)), so that the nucleation rate is

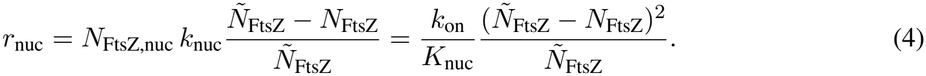

In our simulations we assume that attachment to a nucleation competent monomers happens at the same rate as filament growth, therefore *k*_nuc_ = *k*_on_, leading to the right side of eq.4.

#### 4.8.4 Filament growth

Free monomers attach only to uncapped growing ends, so that the growth rate is

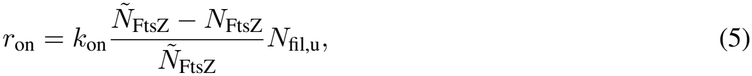

where *k*_on_ is the rate constant of monomer attachment to a single uncapped growing end and *N*_fil,u_ is the number of filaments with an uncapped growing end.

#### 4.8.5 Hydrolysis

Hydrolysis is the only energy-consuming step in the model. FtsZ is GTP-bound when it binds to a filament or to a regulator and hydrolyses its GTP to GDP with the rate constant *k*_hyd_ = 1/*τ*_hyd_, where *τ*_hyd_ is the mean time until hydrolysis of a single GTP-bound FtsZ. We neglect the reverse reaction, as it is rare and does not alter filament behaviour. The hydrolysis rate is

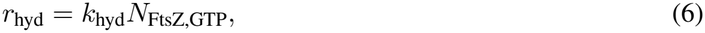

where *N*_FtsZ,GTP_ is the number of GTP-bound FtsZ that are not free monomers.

#### 4.8.6 Filament shrinkage

Monomers detach from the shrinking end, opposite to the growing end, with the rate

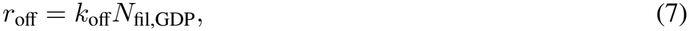

where *k*_off_ is the rate constant of monomer detachment from a single shrinking end and *N*_fil,GDP_ is the number of filaments whose terminal monomer at the shrinking end is GDP-bound; filaments shrink only from a GDP-bound end. Detachment of a monomer from a dimer releases two free monomers (de-nucleation).

#### 4.8.7 Capping

Capping is the binding of a regulator to the growing end of a filament, which blocks further monomer attachment at this end. In analogy to growth, the capping rate is

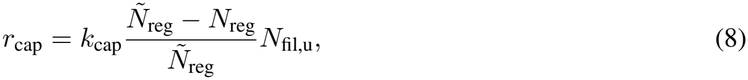

where *k*_cap_ is the rate constant of capping a single uncapped growing end when all regulators are free, *Ñ*_reg_ is the total number of regulators and *N*_reg_ the number of regulators bound to FtsZ. FtsZ L272E caps by binding to the growing end as a monomer would [38]. We therefore use the reference value *k*_cap_ = *k*_on_*Ñ*_reg_/*Ñ*_FtsZ_, so that a free FtsZ L272E molecule binds a growing end with the same rate constant per molecule as a free FtsZ monomer, and *k*_cap_ = *k*_on_ for *Ñ*_reg_ = *Ñ*_FtsZ_.

#### 4.8.8 Sequestration

Sequestration is the process of removing monomeric FtsZ from the system and is modeled as a second-order process. In analogy to nucleation, a free regulator in a sequestration-competent conformation binds a free FtsZ monomer. The number of sequestration-competent regulators is (*Ñ*_reg_ ™ *N*_reg_)/*K*_seq_, which defines the dimensionless sequestration equilibrium constant *K*_seq_ ≥ 1. The sequestration rate is

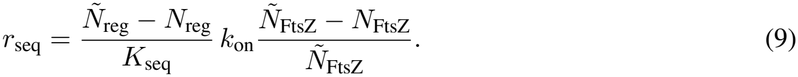

We retain *K*_seq_ in eq. (9) for completeness but set *K*_seq_ = 1 throughout, so that every free regulator can sequester and the effect of sequestration is maximal. A sequestration event is implemented as the formation of a dimer of one regulator and one GTP-bound FtsZ, which counts towards *N*_FtsZ_ and *N*_reg_. The FtsZ in this dimer must hydrolyse its GTP before the dimer can dissociate. Even under this maximal assumption, capping dominates at sub-stoichiometric regulator-to-FtsZ ratios (**fig**. 5).

#### 4.8.9 Severing

Severing breaks a filament into two shorter filaments by cutting the bond between two adjacent subunits. Every internal bond is cut with equal probability, so that the severing rate is

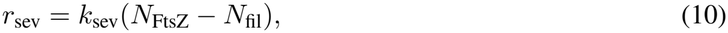

where *k*_sev_ is the severing rate constant per bond and *N*_fil_ is the number of filaments. The factor *N*_FtsZ_ ™*N*_fil_ is the total number of internal bonds, since a filament of *L* subunits contains *L*™1 bonds. Severing is simulated without sequestration, so that all FtsZ counted in *N*_FtsZ_ is polymeric.

To compare severing and capping on equal terms in **fig**. 5, we chose *k*_sev_ such that the severing rate equals the capping rate in the unregulated steady state. We evaluate the capping rate, eq. (8), with no regulator bound, *N*_reg_ = 0, and with all growing ends uncapped, *N*_fil,u_ = *N*_fil_. Using the steady-state averages, eq. (1), of a simulation without regulators, this condition reads

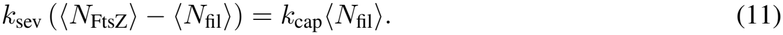

Solving for *k*_sev_ gives

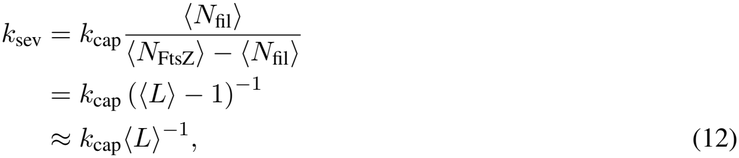

where ⟨*L*⟩ = ⟨*N*_FtsZ_⟩/⟨*N*_fil_⟩ is the mean filament length in the unregulated steady state, and we used ⟨*L*⟩™1 ≈ ⟨*L*⟩ for long filaments.

## 5 Data and code availability

### 5.1 Data availability

Raw fluorescence microscopy images and full datasets supporting the findings of this study will be made publicly available upon publication via the online repository of the Institute of Science and Technology Austria (ISTA). Source data will be provided with this paper.

### 5.2 Code availability

Python scripts used for image analysis and quantification will be made publicly available on GitHub. The compilable source code to our Gillespie solver, a Python interface for analysing its output and scripts for plot generation will be made publicly available via GitHub/OSF.

## 6 Acknowledgements

## 6.1 Acknowledgements

We thank all members of the Loose and Šarić groups at ISTA for helpful discussions and for critical reading of the manuscript. This work was supported by the Scientific Service Units (SSU) of ISTA through resources provided by the Imaging and Optics Facility (IOF), the Lab Support Facility (LSF) and Scientific Computing (SciComp). This work was funded by the Austrian Science Fund (FWF) Principal Investigator Grant PAT1394325, “Mechanisms and evolution of FtsZ regulation” to M.L. and a Wellcome Trust Discovery Award to A.S..

## 6.2 Author contributions

X.Y. and M.L. conceived the study. X.Y. designed and performed the experiments and analysed the data. M.L. developed the research strategy, analysed the data and the structure predictions with help from X.Y. F.W. developed the model under the guidance of A.S. and M.L., A.S. supervised the modelling work and contributed critical input on its development. M.L. and A.S. supervised the project and acquired funding. X.Y. and M.L. wrote the manuscript, with F.W. and A.S. writing the sections on the model. All authors reviewed and edited the manuscript.

## S1 Supplementary figures

**Figure S1:**
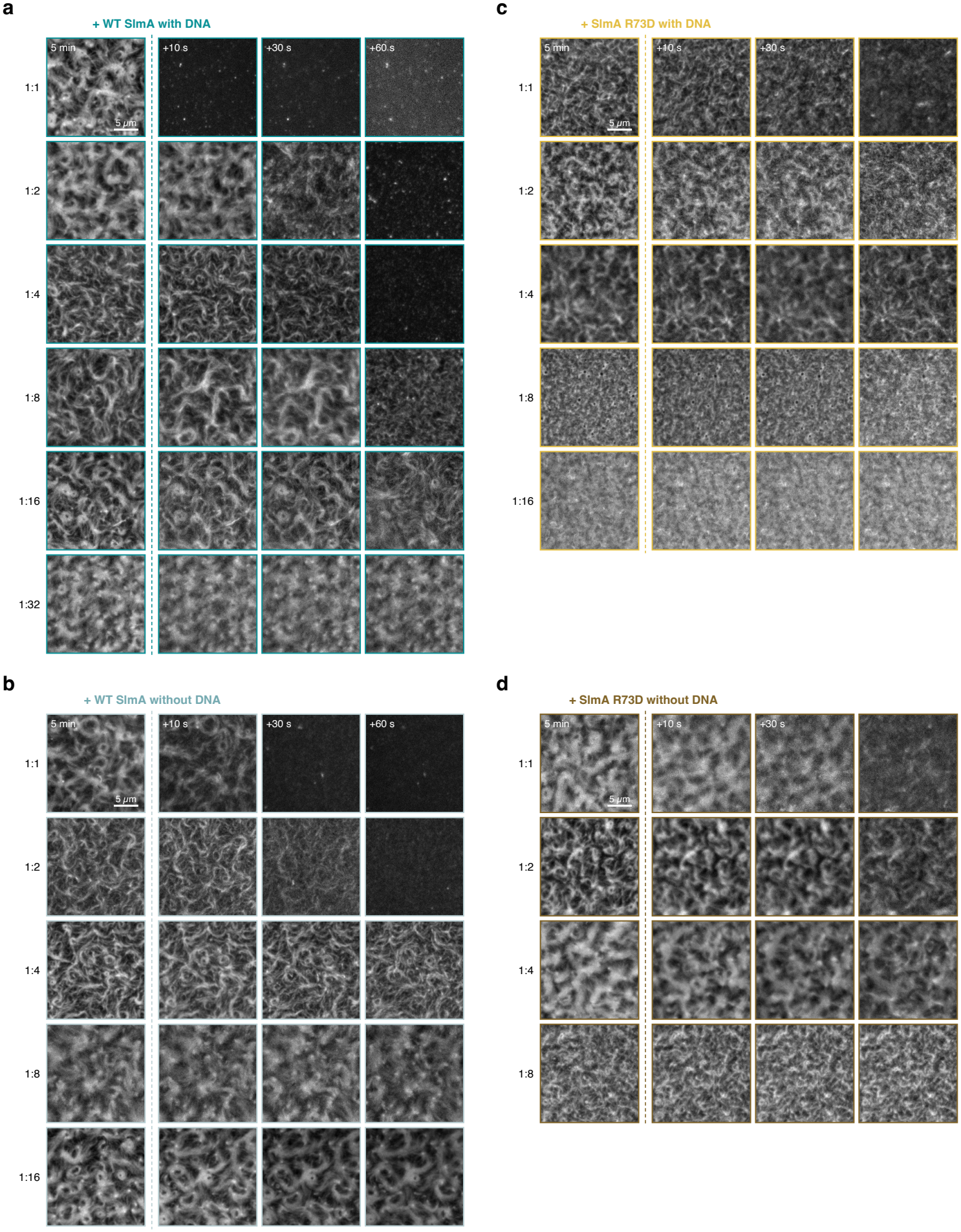
FtsZ filament disassembly by wild-type SlmA and SlmA R73D, with and without DNA. TIRF micrographs of FtsZ filament networks on FtsA-containing supported lipid bilayers (1.25 µM FtsZ) before and after addition of SlmA at the indicated SlmA:FtsZ molar ratios. **a**, Wild-type SlmA with DNA (1:1 to 1:32). **b**, Wild-type SlmA without DNA (1:1 to 1:16). **c**, SlmA R73D with DNA (1:1 to 1:16). **d**, SlmA R73D without DNA (1:1 to 1:8). For each condition, the first column shows the FtsZ network 5 min after the start of the experiment, immediately before SlmA addition (dashed line). The second and third columns show the same field 10 s and 30 s after SlmA addition. The last column shows the frame 60 s after addition. Image borders are colour-coded as in Fig. 1. Scale bar, 5 µm.

**Figure S2:**
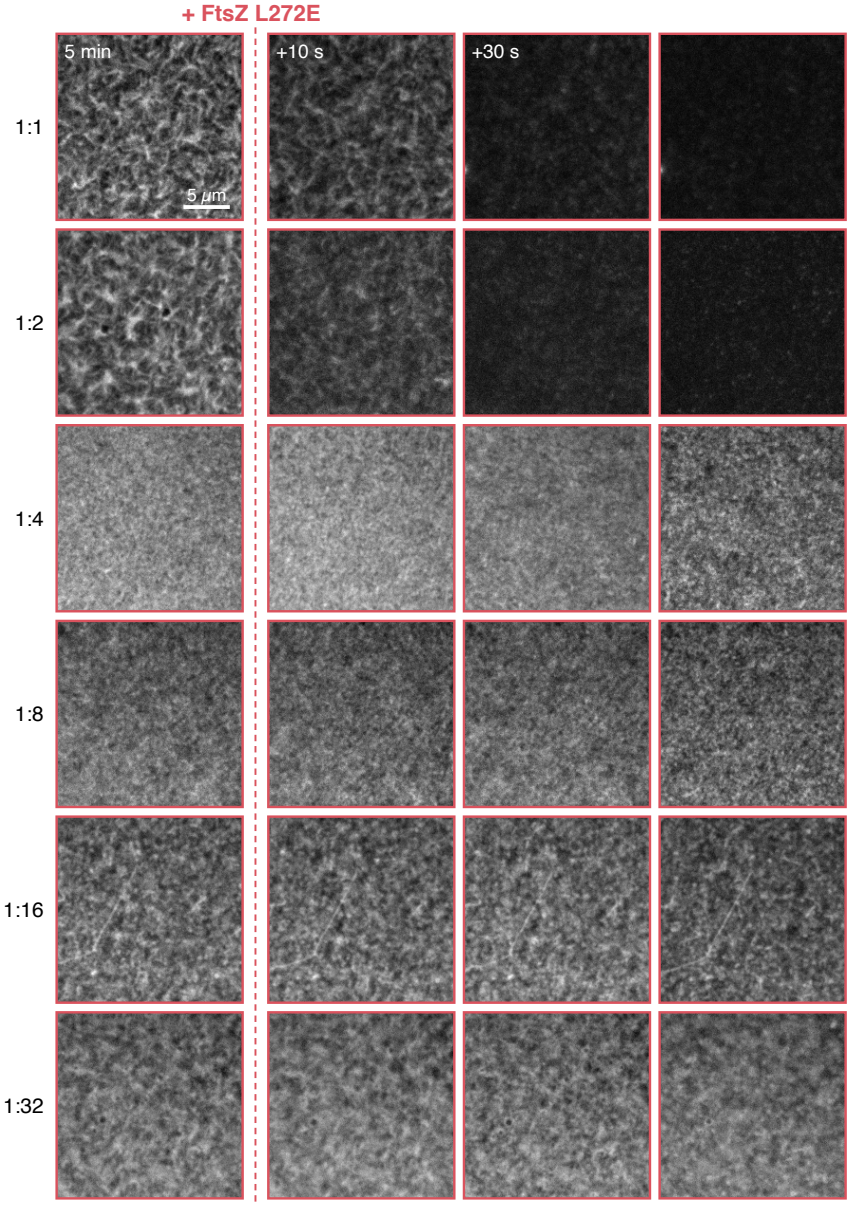
FtsZ L272E-induced disassembly of FtsZ filaments. TIRF micrographs of FtsZ filament networks on FtsA-containing supported lipid bilayers (1.25 µM FtsZ ) before and after addition of FtsZ L272E at FtsZ:FtsZ L272E molar ratios from 1:1 to 1:32. The first column shows the FtsZ network 5 min after the start of the experiment, immediately before FtsZ L272E addition (dashed line). The second and third columns show the same field 10 s and 30 s after FtsZ L272E addition. Scale bar, 5 µm.

**Figure S3:**
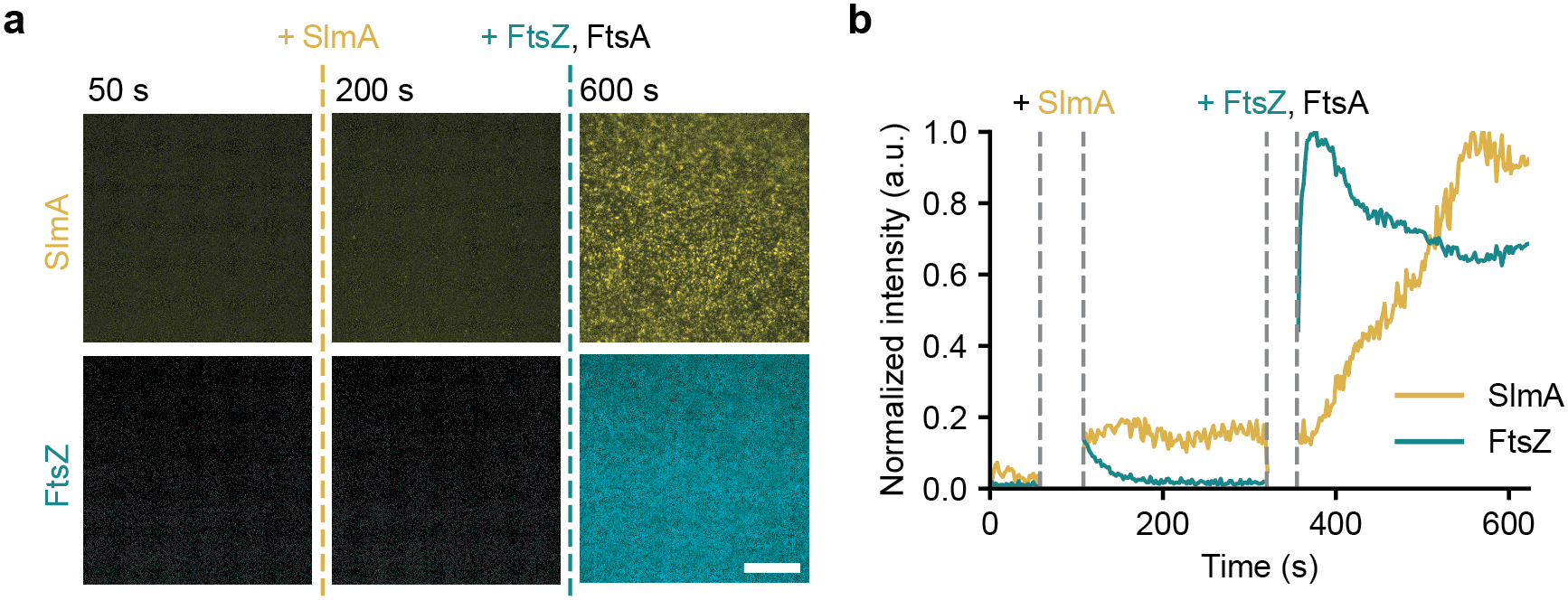
SlmA does not bind to membranes: **(a)** TIRF fluorescence micrographs of SlmA and FtsZ. In the absence of FtsZ, no Cy5-labeled SlmA (78 nM) was detected on the membrane surface. Only after the addition of Alexa488-labeled FtsZ (1.25 *µ*M, 5 mM GMPCPP and unlabeled FtsA (0.4 *µ*M) did the fluorescence intensity corresponding to SlmA increase. **(b)** Intensity curves corresponding to the micrographs shown in panel (a). The segments between the gray dashed lines indicate the time points at which proteins were added.

**Figure S4:**
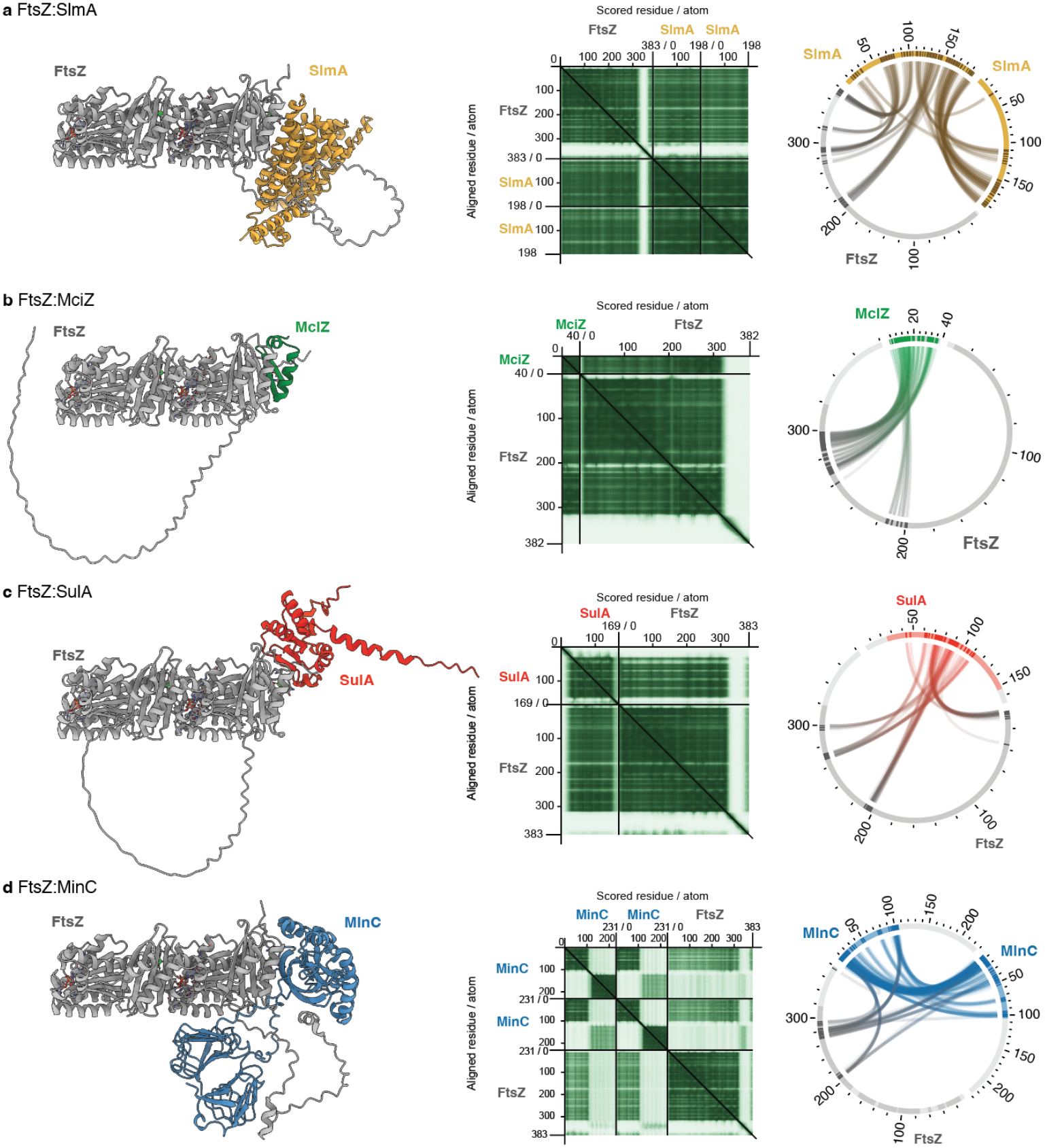
AlphaFold3 predictions of complexes between FtsZ and four known FtsZ-binding proteins: **(a)** SlmA (yellow), **(b)** MciZ (green), **(c)** SulA (red), and **(d)** MinC (blue). For each complex, the predicted structure of the FtsZ-binding protein bound to a single FtsZ subunit (grey) was aligned with the terminal subunit of an FtsZ filament (PDB 3VOA, grey) to visualize the predicted binding mode in the context of a filament end. Next to each predicted complex structure, the predicted aligned error (PAE) map for the corresponding AlphaFold3 model is shown, with the FtsZ and binding-partner axes labeled and residue numbers indicated. To the right of each PAE map, a circular contact map shows the predicted interchain contacts between FtsZ (gray) and the respective binding protein (colored arcs), with connecting lines indicating residue pairs predicted to be in contact according to the cLIR score [57].

**Figure S5:**
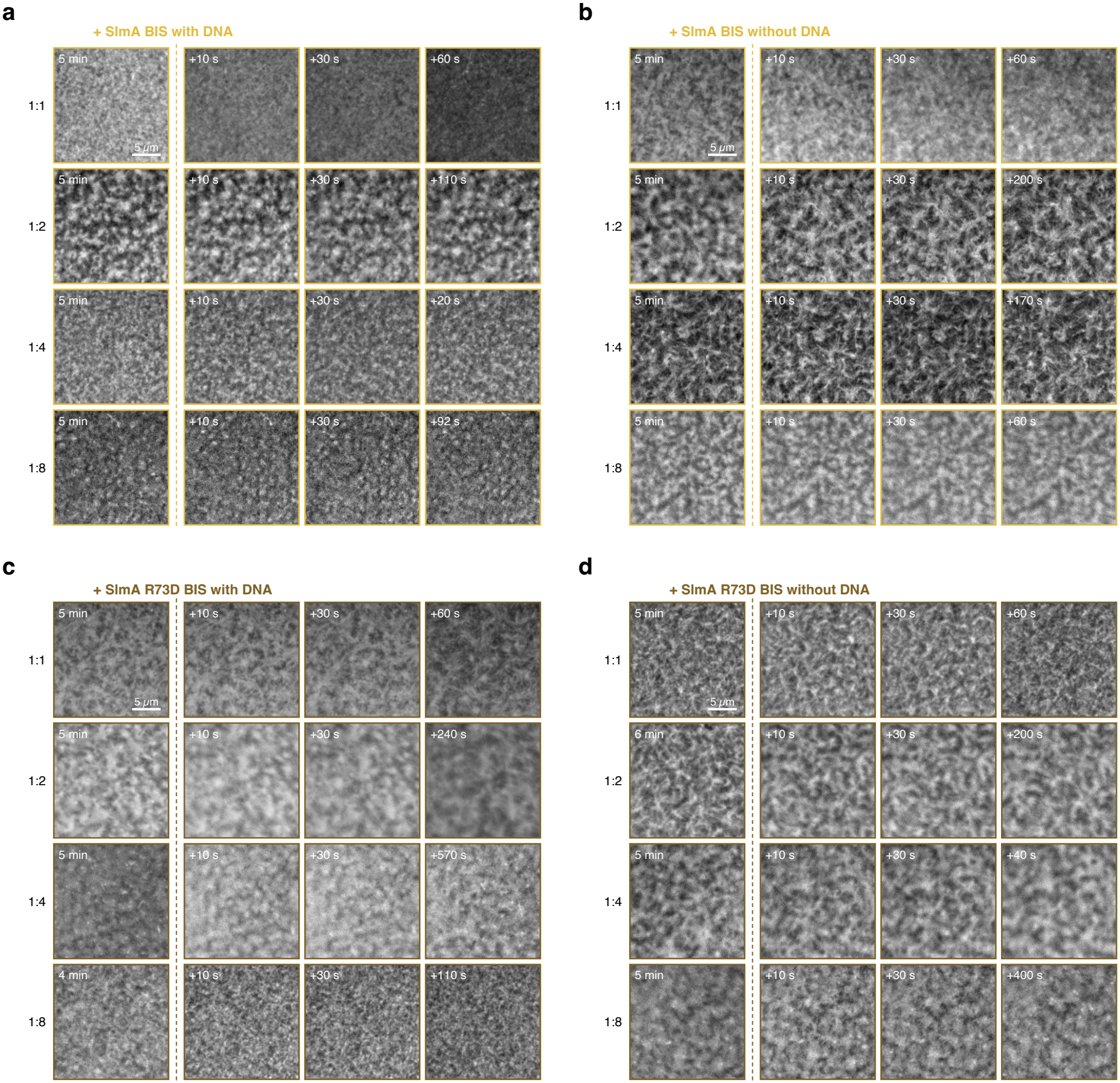
FtsZ filament disassembly by SlmA BIS and SlmA R73D BIS mutants, with and without DNA. TIRF micrographs of FtsZ filament networks on FtsA-containing supported lipid bilayers (1.25 µM FtsZ) before and after addition of SlmA mutants at SlmA:FtsZ molar ratios from 1:1 to 1:8. **a**, SlmA BIS with DNA. **b**, SlmA BIS without DNA. **c**, SlmA R73D BIS with DNA. **d**, SlmA R73D BIS without DNA. For each row, the first column shows the FtsZ network before SlmA addition (dashed line). The second and third columns show the same field 10 s and 30 s after addition. The last column shows the frame of lowest FtsZ intensity. Because the time to reach lowest intensity differed between conditions, the time of each frame is indicated in the image. Scale bar, 5 µm.

**Figure S6:**
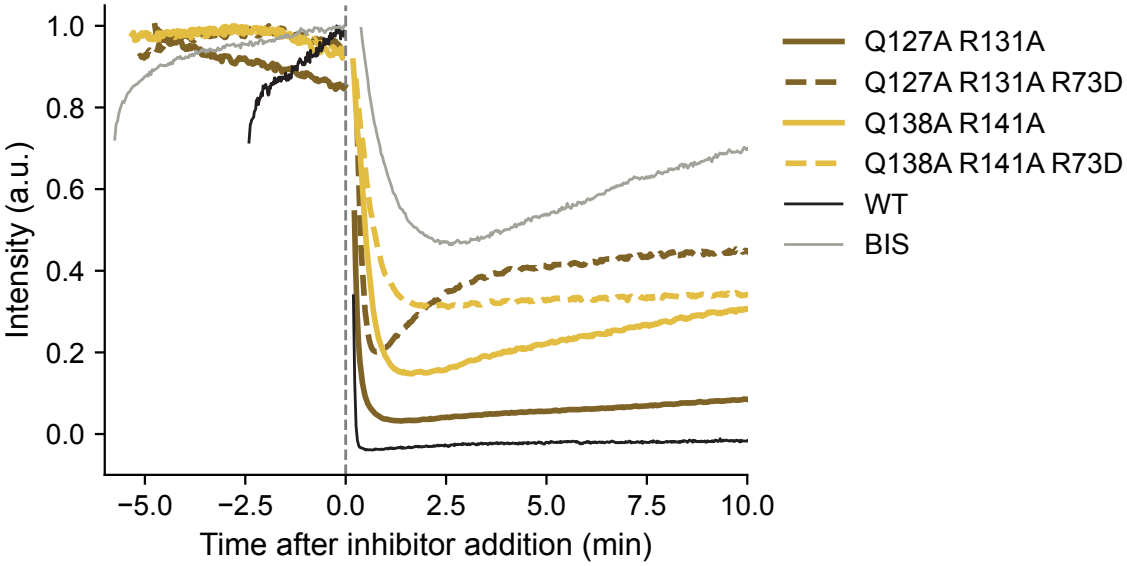
SlmA double mutants are still active: Fluorescence intensity curves showing depolymerization of FtsZ filaments after addition of 1.25 *µ*M SlmA Q127A R131A (brown), SlmA Q138A R141A (yellow), WT SlmA (black) or the quadruple mutant SlmA BIS (grey) with FtsZ = 1.25 *µ*M, FtsA = 0.4 *µ*M. Dashed lines show the corresponding triple mutants carrying the additional R73D mutation. Each trace was normalized individually, setting its maximum intensity before SlmA addition to 1 and the background intensity after EDTA addition to 0 (not shown). Traces are aligned to the time of SlmA addition (*t* = 0). SlmA Q127A R131A retains wildtype activity, while the activity of SlmA Q138A R141A is slightly decreased. Both double mutants are more active than the quadruple BIS mutant and the double mutants additionally carrying R73D.

## S2 Supplementary movies

Supplementary Movie 1: **Depolymerization of FtsZ filaments by wild-type SlmA with DNA**. TIRF movie of treadmilling FtsZ filaments on supported lipid bilayers (1.25 µM FtsZ, 0.4 µM FtsA) before and after addition of SBS-bound wild-type SlmA (1.25 µM). Left, intensity-normalized; right, raw fluorescence intensities. Corresponds to Fig. 1c,e. Scale bar, 10 µm.

Supplementary Movie 2: **Depolymerization of FtsZ filaments by SlmA R73D with DNA**. As Supplementary Movie 1, but after addition of SBS-bound SlmA R73D (1.25 µM). Left, intensity-normalized; right, raw fluorescence intensities. Corresponds to Fig. 1d,f. Scale bar, 10 µm.

Supplementary Movie 3: **SlmA is recruited to the membrane only by FtsZ**. TIRF movie of single Cy5-labeled SlmA molecules (78 nM) on supported lipid bilayers, first alone and then after addition of Alexa488-labeled FtsZ (1.25 µM) and unlabeled FtsA (0.4 µM) with 1 mM GMPCPP. Left, SlmA; middle, FtsZ; right, merge. Corresponds to Fig. S3. Scale bar, 10 µm.

Supplementary Movie 4: **Single-molecule imaging of SlmA on treadmilling FtsZ filaments**. TIRF movie of single Cy5-labeled wild-type SlmA molecules binding to membrane-bound FtsZ filaments in the presence of FtsA; only the SlmA channel is shown. Corresponds to Fig. 2g. Scale bar, 10 µm.

Supplementary Movie 5: **Reduced depolymerization of FtsZ filaments by SlmA BIS with DNA**. As Supplementary Movie 1, but after addition of SBS-bound SlmA BIS (Q127A R131A Q138A R141A; 1.25 µM). Left, intensity-normalized; right, raw fluorescence intensities. Corresponds to Fig. 3e. Scale bar, 10 µm.

Supplementary Movie 6: **SlmA R73D BIS barely affects FtsZ filaments**. As Supplementary Movie 1, but after addition of SBS-bound SlmA R73D BIS (1.25 µM). Left, intensity-normalized; right, raw fluorescence intensities. Corresponds to Fig. 3e. Scale bar, 10 µm.

Supplementary Movie 7: **Stochastic simulation of treadmilling FtsZ filaments**. Gillespie simulation of treadmilling filaments coupled to a finite pool of monomers and regulators (**sec**. 4.8). This simulation considers (de-)nucleation, growth, shrinkage, hydrolysis and filament capping through regulators. **a)** Visual representation of the simulation as it is stored in the implementation of the Gillespie solver. Top: Fractions of free FtsZ monomers, (*Ñ*_FtsZ_ ™ *N*_FtsZ_)/*Ñ*_FtsZ_, and of free regulators, (*Ñ*_reg_ ™ *N*_reg_)/*Ñ*_reg_, overlaid with the total numbers *Ñ*_FtsZ_ and *Ñ*_reg_. Bottom: GTP-bound FtsZ in dark cyan, GDP-bound FtsZ in light cyan, and regulatory proteins in yellow, filaments grow from left to right, new filaments nucleate appear at from the top. Time is displayed in units of 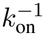. **b)** Current time between events, in units of 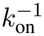, showing the current timestep (Δ*t*), nucleation (nuc), de-nucleation (denuc), growth (on), shrinkage (off) and hydrolysis (hyd). Once the system has reached steady state, nucleation and de-nucleation occur at equal rates, as do growth, shrinkage and hydrolysis. **c)** Filament length distributions, the current distribution is shown as grey bars, the averaged distribution until the current time *t* is shown as a cyan, continuous line.

## S3 Supplementary text

### S3.1 Simulation parameters and observables

**Table S1:**
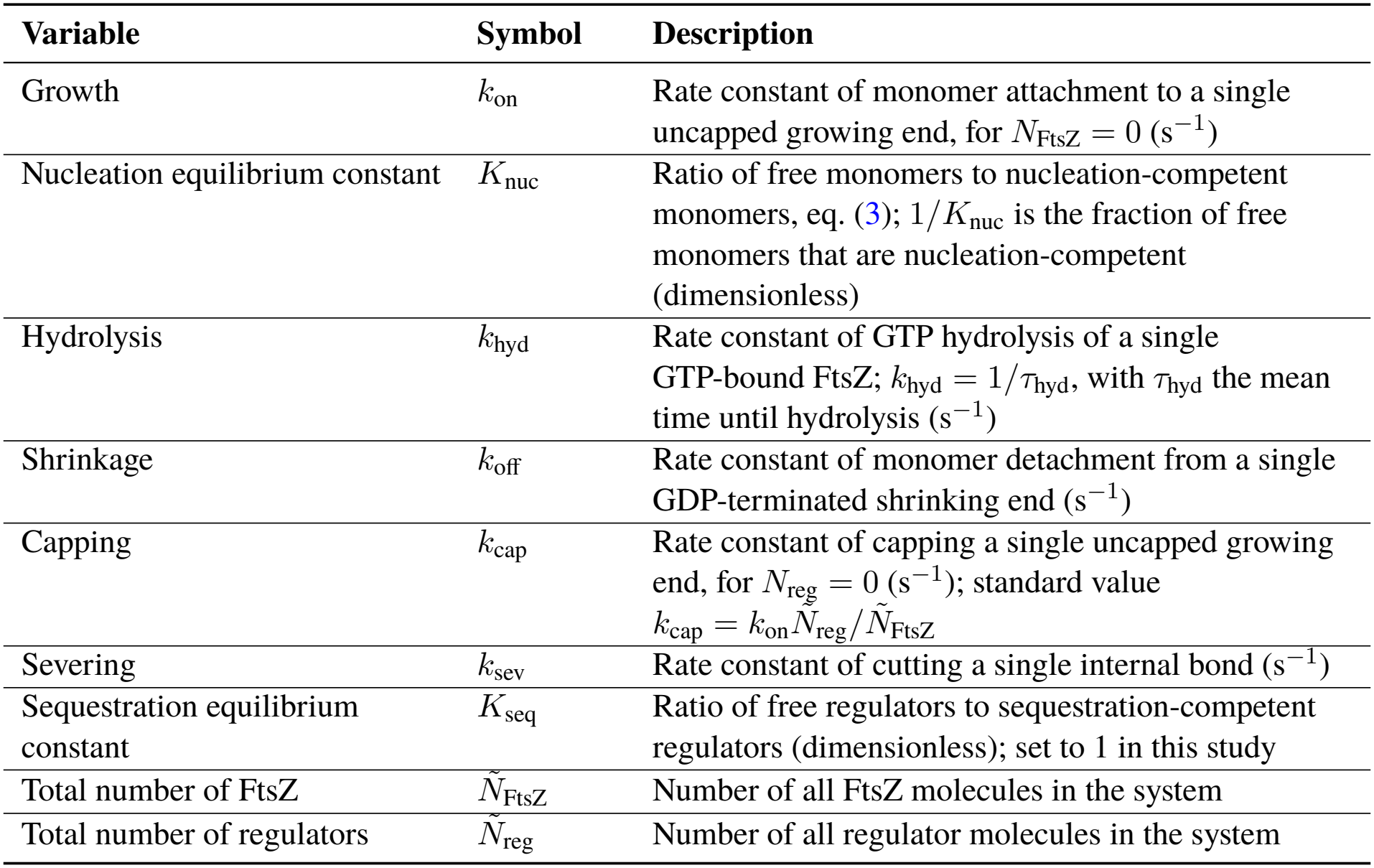
Simulation parameters. Rate constants and molecule numbers that serve as input to the filament simulations.

| **Variable** | **Symbol** | **Description** |
|---|---|---|
| Growth | *k*<sub>on</sub> | Rate constant of monomer attachment to a single uncapped growing end, for *N*<sub>FtsZ</sub> = 0 (s<sup>−1</sup>) |
| Nucleation equilibrium constant | *K*<sub>nuc</sub> | Ratio of free monomers to nucleation-competent monomers, eq. (3); 1/*K*<sub>nuc</sub> is the fraction of free monomers that are nucleation-competent (dimensionless) |
| Hydrolysis | *k*<sub>hyd</sub> | Rate constant of GTP hydrolysis of a single GTP-bound FtsZ; *k*<sub>hyd</sub> = 1/*τ*<sub>hyd</sub>, with *τ*<sub>hyd</sub> the mean time until hydrolysis (s<sup>−1</sup>) |
| Shrinkage | *k*<sub>off</sub> | Rate constant of monomer detachment from a single GDP-terminated shrinking end (s<sup>−1</sup>) |
| Capping | *k*<sub>cap</sub> | Rate constant of capping a single uncapped growing end, for *N*<sub>reg</sub> = 0 (s<sup>−1</sup>); standard value $k_{\mathrm{cap}} = k_{\mathrm{on}}\tilde{N}_{\mathrm{reg}}/\tilde{N}_{\mathrm{FtsZ}}$ |
| Severing | *k*<sub>sev</sub> | Rate constant of cutting a single internal bond (s<sup>−1</sup>) |
| Sequestration equilibrium constant | *K*<sub>seq</sub> | Ratio of free regulators to sequestration-competent regulators (dimensionless); set to 1 in this study |
| Total number of FtsZ | $\tilde{N}_{\mathrm{FtsZ}}$ | Number of all FtsZ molecules in the system |
| Total number of regulators | $\tilde{N}_{\mathrm{reg}}$ | Number of all regulator molecules in the system |

**Table S2:** Rates of all processes in the Gillespie scheme. Each rate *r*_*i*_ is calculated from the rate constants and molecule numbers of Table S1 and the state variables of Table S3. *Ñ*_FtsZ_ ™ *N*_FtsZ_ and *Ñ*_reg_ ™ *N*_reg_ are the numbers of free FtsZ monomers and free regulators. With the total rate *R* = ∑_*i*_ *r*_*i*_, the Gillespie solver draws the next timestep from Δ*t* = ™ ln(*u*_1_)/*R*, which is exponentially distributed with mean 1/*R*, and executes process *i* with probability *r*_*i*_/*R*.

| Process | Rate | Requires | Eq. |
|---|---|---|---|
| Nucleation | $r_{\mathrm{nuc}} = \frac{k_{\mathrm{on}}}{K_{\mathrm{nuc}}} \frac{\left(\tilde{N}_{\mathrm{FtsZ}} - N_{\mathrm{FtsZ}}\right)^2}{\tilde{N}_{\mathrm{FtsZ}}}$ | A nucleation-competent free monomer (fraction 1/*K*<sub>nuc</sub> of free monomers) and a second free monomer | (4) |
| Growth | $r_{\mathrm{on}} = k_{\mathrm{on}} \frac{\tilde{N}_{\mathrm{FtsZ}} - N_{\mathrm{FtsZ}}}{\tilde{N}_{\mathrm{FtsZ}}} N_{\mathrm{fil,u}}$ | A free monomer and an uncapped growing end | (5) |
| Hydrolysis | *r*<sub>hyd</sub> = *k*<sub>hyd</sub>*N*<sub>FtsZ,GTP</sub> | A GTP-bound FtsZ in a filament or bound to a regulator; irreversible | (6) |
| Shrinkage | *r*<sub>off</sub> = *k*<sub>off</sub>*N*<sub>fil,GDP</sub> | A GDP-bound terminal monomer at the shrinking end; de-nucleates the filament if it is a dimer | (7) |
| Capping | $r_{\mathrm{cap}} = k_{\mathrm{cap}} \frac{\tilde{N}_{\mathrm{reg}} - N_{\mathrm{reg}}}{\tilde{N}_{\mathrm{reg}}} N_{\mathrm{fil,u}}$ | A free regulator and an uncapped growing end; blocks further growth at that end | (8) |
| Sequestration | $r_{\mathrm{seq}} = \frac{\tilde{N}_{\mathrm{reg}} - N_{\mathrm{reg}}}{K_{\mathrm{seq}}} k_{\mathrm{on}} \frac{\tilde{N}_{\mathrm{FtsZ}} - N_{\mathrm{FtsZ}}}{\tilde{N}_{\mathrm{FtsZ}}}$ | A sequestration-competent free regulator (fraction 1/*K*<sub>seq</sub>, set to 1 here) and a free monomer | (9) |
| Severing | *r*<sub>sev</sub> = *k*<sub>sev</sub>(*N*<sub>FtsZ</sub> − *N*<sub>fil</sub>) | An internal bond; a filament of *L* subunits contains *L* − 1 of them | (10) |

**Table S3:**
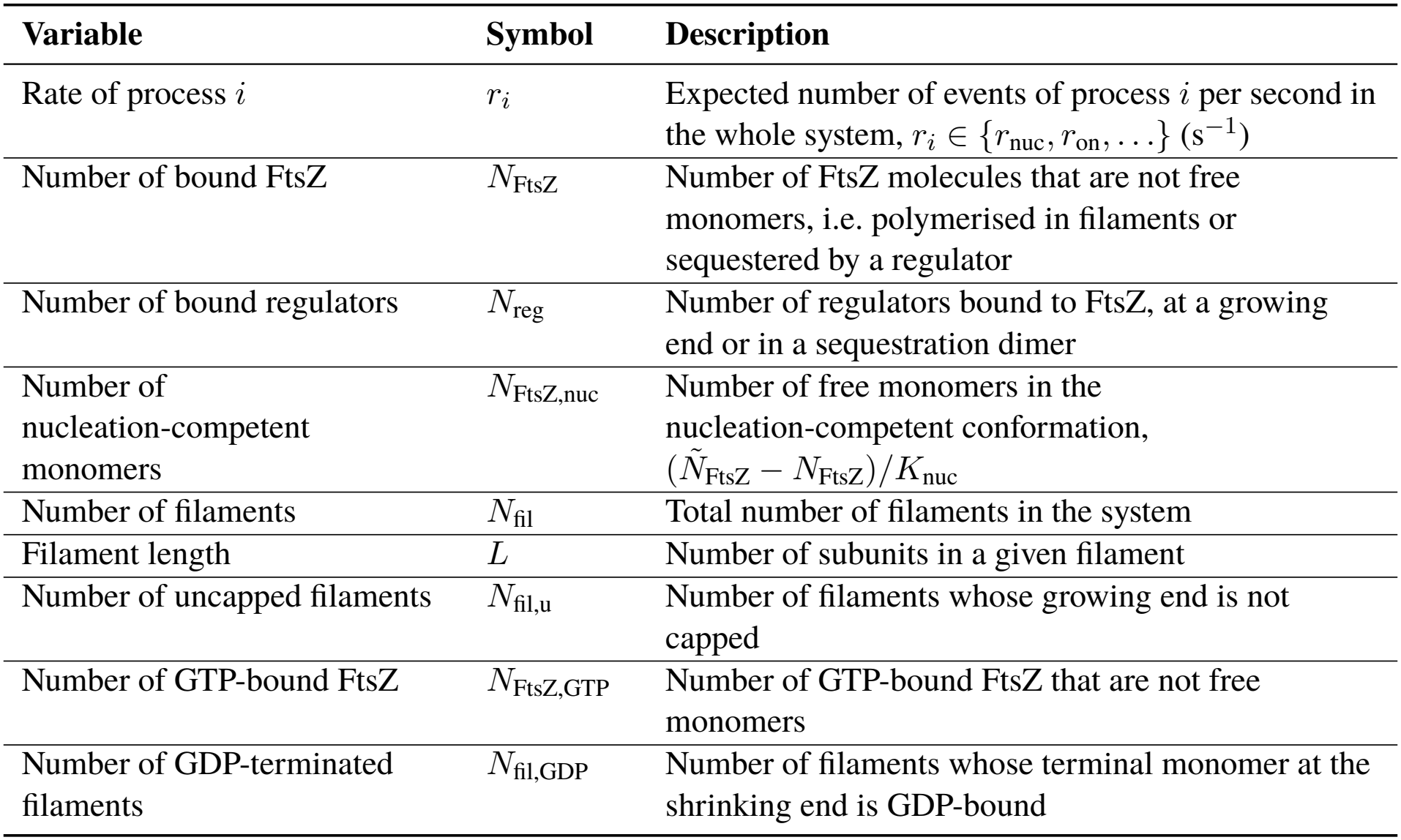
State variables. Quantities that describe the current state of the simulation and enter the rates of Table S2. ⟨*X*⟩ denotes the steady-state time average of a state variable *X*, eq. (1).

| **Variable** | **Symbol** | **Description** |
|---|---|---|
| Rate of process *i* | *r*<sub>*i*</sub> | Expected number of events of process *i* per second in the whole system, *r*<sub>*i*</sub> ∈ {*r*<sub>nuc</sub>, *r*<sub>on</sub>, . . .} (s<sup>−1</sup>) |
| Number of bound FtsZ | *N*<sub>FtsZ</sub> | Number of FtsZ molecules that are not free monomers, i.e. polymerised in filaments or sequestered by a regulator |
| Number of bound regulators | *N*<sub>reg</sub> | Number of regulators bound to FtsZ, at a growing end or in a sequestration dimer |
| Number of nucleation-competent monomers | *N*<sub>FtsZ,nuc</sub> | Number of free monomers in the nucleation-competent conformation, (*Ñ*<sub>FtsZ</sub> − *N*<sub>FtsZ</sub>)/*K*<sub>nuc</sub> |
| Number of filaments | *N*<sub>fil</sub> | Total number of filaments in the system |
| Filament length | *L* | Number of subunits in a given filament |
| Number of uncapped filaments | *N*<sub>fil,u</sub> | Number of filaments whose growing end is not capped |
| Number of GTP-bound FtsZ | *N*<sub>FtsZ,GTP</sub> | Number of GTP-bound FtsZ that are not free monomers |
| Number of GDP-terminated filaments | *N*<sub>fil,GDP</sub> | Number of filaments whose terminal monomer at the shrinking end is GDP-bound |

### S3.2 Choice of experimental observables

**Figure S7:**
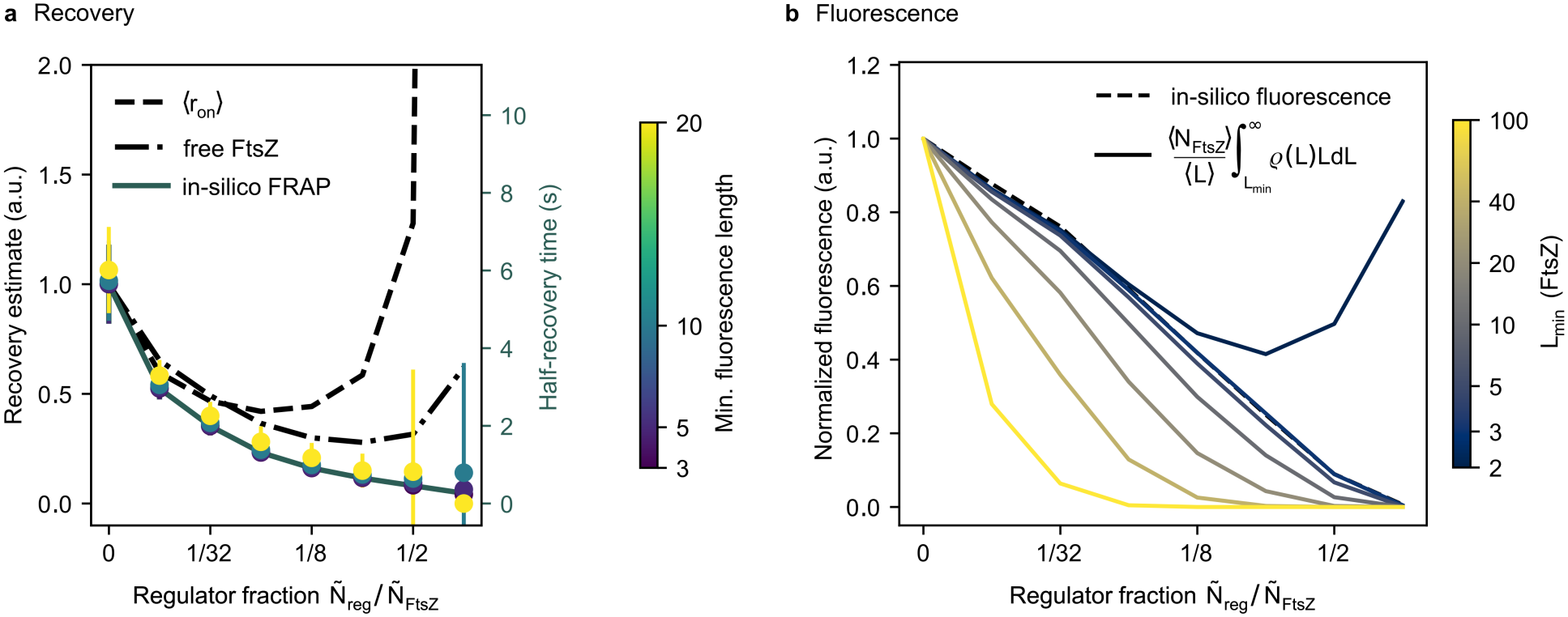
*In silico* observables for comparison to experimental data. **a)** Candidates for relative half-recovery time. Coloured curves show half-recovery times measured by *in silico* FRAP, considering only filaments of at least the minimum fluorescence length (right axis); black lines show estimates of the recovery time based on system properties (left axis). The system properties used as proxies for recovery times are the total growth rate ⟨*r*_on_⟩, eq. (5), and the fraction of free monomers 1 ™ ⟨*N*_FtsZ_⟩/*Ñ*_FtsZ_. **b)** Candidates for fluorescence measurements. *In silico* fluorescence measurements only consider fluorescence from FtsZ that is not a free monomer. Alternatively, the number of FtsZ in filaments of length *L* ≥ *L*_min_, eq. (S1), discounts short filaments and sequestered FtsZ.

Since it was not immediately clear how to map experimentally determined half-recovery times and fluorescence to the simulated values determined by our Gillespie solver, we introduced different simulated observables to represent them. A straight-forward approach are *in silico* fluorescence experiments, whereby each monomer that was added into the simulated system received a marker. A marker sitting on a monomer is counted as one fluorescent signal and stays active until the monomer detaches from the filament again or there is a bleaching event which deactivates it. The total number of markers at a given point in time was used as a proxy for fluorescence, depicted as *in silico* fluorescence in **fig**. S7**b**. Furthermore, by setting all markers to be non-fluorescent at pre-determined points within the simulation, an *in silico* FRAP experiment could be run, where the half-recovery time was determined by the time it takes to recover half the amount of fluorescent markers in the system.

Experimentally measured fluorescence both in FRAP and total fluorescence could also originate mostly from long filaments, as short filaments could be disregarded by noise filters. We therefore additionally implemented a version of *in silico* FRAP which only considers fluorescence from filaments above a certain length, but found that even when disregarding filaments of up to 20 monomers the results for half-recovery times were consistent with considering all fluorescent markers in the system (**fig**. S7**a**). However, we found a large difference between the recovery estimated from the total growth rate ⟨*r*_on_⟩, eq. (5), and from the fraction of free monomers 1 ™ ⟨*N*_FtsZ_⟩/*Ñ*_FtsZ_. These observables proved to be relevant when trying to gauge the recovery of fluorescence of long polymers. At high regulator fractions, we expected the system to contain many short filaments which may not fluoresce in experiments, but which contributed to recovery in our *in silico* experiments. We therefore used ⟨*r*_on_⟩ as a proxy for the relative half-recovery times in **fig**. 5.

We also found a different response to regulators when measuring the total fluorescence of simulated systems of treadmilling FtsZ filaments. Instead of counting fluorescent markers, we integrate the normalised filament length distribution *ρ*(*L*) from a lower limit *L*_min_. The proxy for fluorescence is then the number of FtsZ in filaments with *L* ≥ *L*_min_,

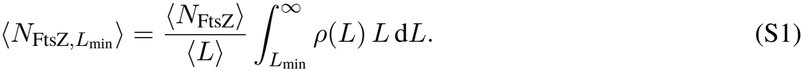

**Fig**. S7**b** shows that there is a difference between *in silico* fluorescence, which excludes sequestered monomers and fluorescence based on eq (S1). We assume that filaments short filaments bind the membrane only transiently and are therefore not detected in TIRF experiments. We therefore used *L*_min_ = 5 for the relative fluorescence in **fig**. 5.

### S3.3 Normalised intensity from fluorescence measurements

**Figure S8:**
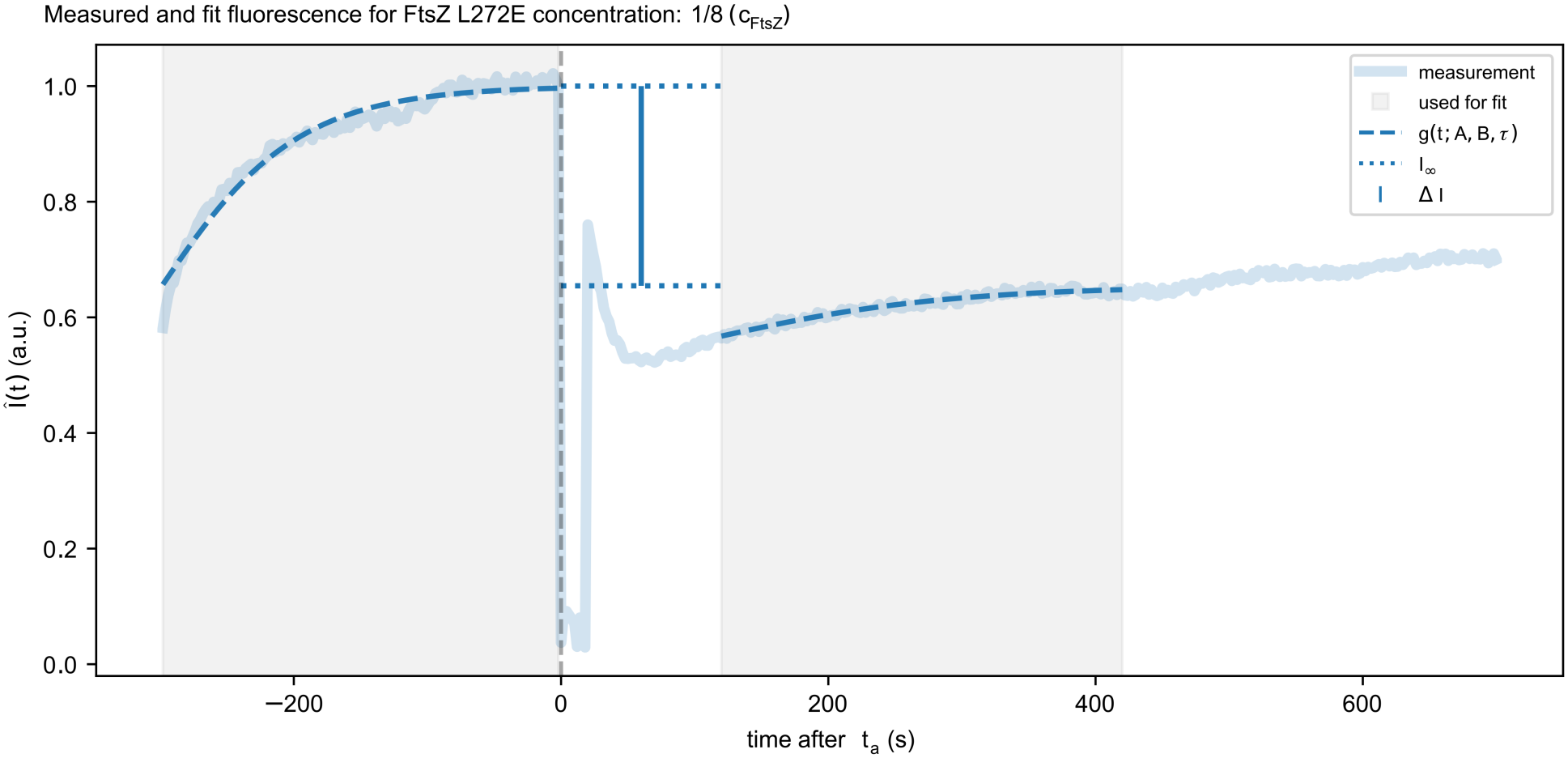
Visualisation of extracting the normalised decrease in fluorescence intensity from experimentally determined data. The semi-transparent line shows the fluorescence intensity of one replicate with 0.15625 µM FtsZ L272E and 1.25 µM FtsZ, i.e. a regulator ratio [FtsZ L272E]/[FtsZ] = 1/8. The intensity is normalised to 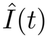. Grey areas mark the fit windows before and after regulator addition at *t*_a_, dashed lines show the fits of eq. (S2), dotted lines the resulting asymptotes 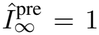 and 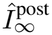, and the vertical line the normalised decrease 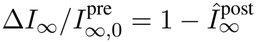. This replicate was not flushed with EDTA, so *I*_bg_ = 0.

We used an automated procedure to extract the normalised decrease in fluorescence from measured fluorescence intensity traces *I*(*t*). Each measurement has two or three stages. (i) equilibration, during which the fluorescence of the surface-bound filaments approached a steady state; (ii) addition of the regulator (FtsZ L272E, or SlmA variants) at time *t*_a_, after which the fluorescence relaxed to a new, lower steady state; and (iii) for some replicates, a final flush with EDTA removed the filaments from the surface, so that only the background fluorescence remained. We aim to find the relative difference between the steady-state fluorescence intensities before (i) and after (ii) the addition of regulators, with respect to the background fluorescence *I*_bg_

Addition of the regulators and adjusting the probe causes a dip in measured fluorescence and we can detect the addition by finding the first point in the trace where fluorescence intensity drops below 20% of the maximum value within the first half of a trace. For traces with a final flush, the background fluorescence intensity *I*_bg_ is the mean of the last 10 data points. For all other traces we set *I*_bg_ = 0.

Not all measured fluorescence intensity curves reached a steady state, as doing so would have required unreasonably long measurement times for each experiment. We therefore estimated the steady state fluorescence intensity *I*_*∞*_ by fitting a Gompertz function,

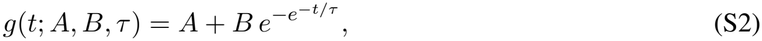

to *I*(*t*) using nonlinear least squares (`scipy.optimize.curve_fit` [76]) and extracting *I*_*∞*_ = lim_*t→∞*_ *g* = *A* + *B*.

We fit Eq. (S2) to the background-subtracted trace *I*_0_(*t*) = *I*(*t*) ™ *I*_bg_ over the equilibration stage, from *t* = 0 to *t*_a_, yielding the expected steady-state fluorescence intensity before adding the regulator, 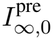. We then normalised the intensity to

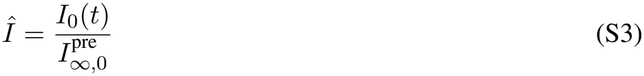

and again fit Eq. (S2) to the window *t*_a_ + 120 s *< t < t*_a_ + 420 s. The asymptote of this fit, 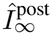 is the normalised steady-state fluorescence in the presence of the regulator.

For each replicate, the normalised fluorescence decrease is defined as

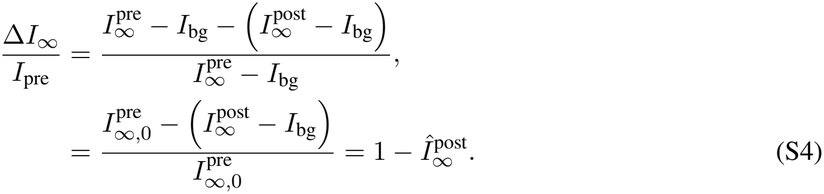

**Fig**. S8 visualises this automated procedure for one replicate at [FtsZ L272E]/[FtsZ] = 1/8.

### S3.4 Determining treadmilling parameters from in vitro data

**Figure S9:**
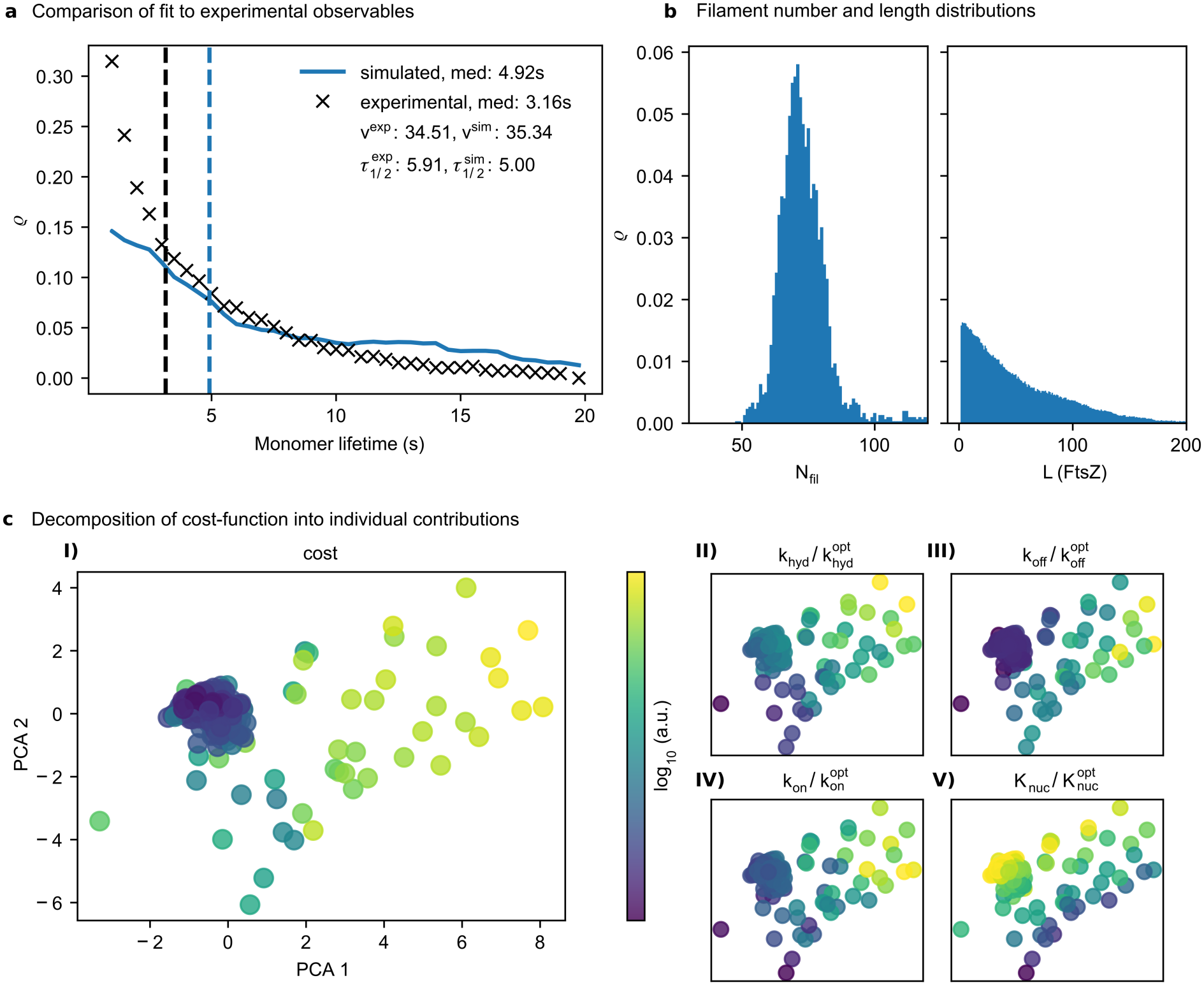
Results of determining treadmilling parameters using Bayesian optimisation. **a)** Comparison of *in vitro* observables [8, 51, 53, 52] to best fit simulation. **b)** Distribution of number of filaments and filament length, connecting exponentially distributed filament lengths to monomer lifetimes. **c)** PCA of log difference vectors of the parameters for each step during the optimisation. The input of the PCA is built from the four-dimensional vector 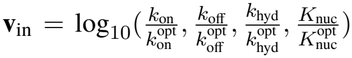. The constructed PCA is colored by the total optimisation cost **(I)** and by each of the entries for **v**_in_ **(II-V)** to gauge the importance of optimising the respective entries in decreasing the optimisation cost.

Extracting the rate constant *k*_on_ from steady-state measurements of the growth rate ⟨*r*_on_⟩ in treadmilling systems with a known steady-state fraction of free monomers, 1 ™ ⟨*N*_FtsZ_⟩/*Ñ*_FtsZ_, is not straightforward. In experimental systems, treadmilling filaments block each other’s paths, a process that has been shown to generate nematic alignment in treadmilling filament systems [62]. Therefore, the experimentally determined growth rate per filament, ⟨*r*_on_⟩/⟨*N*_fil_⟩, depends on the probability *p*_free_(⟨*σ*⟩) that a growing end is not blocked by another filament,

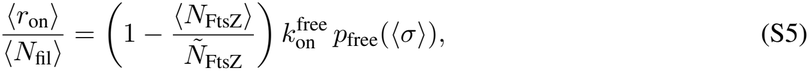

where 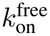 is the rate constant of monomer attachment to an unobstructed growing end and *σ* is the surface density of FtsZ. Our simulation does not resolve space, so that in the unregulated system, where *N*_fil,u_ = *N*_fil_, eq. (5) gives

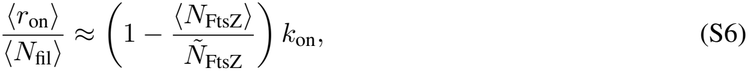

neglecting correlations between the fluctuations of *N*_FtsZ_ and *N*_fil_. The rate constant *k*_on_ of a Gillespie simulation therefore corresponds to 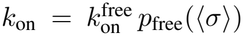, which depends on the surface density ⟨*σ*⟩ of the experiment. The probability *p*_free_ is not necessarily the same in different systems, and its dependence on ⟨*σ*⟩ also couples *k*_on_ to the nematic order of the system, which itself changes with the fraction of free monomers [62].

We show in **fig**. S10**a** that using a previously determined [77] effective hydrolysis rate constant *k*_hyd_ = 0.137 s^*™*1^ constrains the system such that there is no parameter range which matches experimentally determined velocities *v*^exp^, FRAP half-recovery times 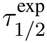 as well as monomer lifetimes *ρ*^exp^(*t*) [8, 51, 53, 52]. We found a better match to experimentally determined values when increasing the hydrolysis rate constant *k*_hyd_ **fig**. S10**b**.

We therefore applied Bayesian optimisation when exploring the parameter space (**fig** S9) to systematically base our treadmilling parameters in **fig**. 5 on experimentally measured behaviour. To obtain the fit to experimentally determined *v*^exp^, 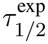 and *ρ*^exp^(*t*) in **fig**. S9**a** we used

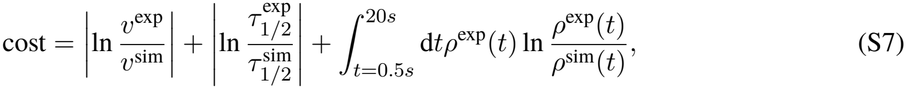

where the limits in 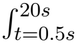 are determined by the experimentally available data. The optimal values according to the Bayesian optimisation scheme are given in Table S4, yielding a Gaussian distribution of the number of filaments *N*_fil_ and an exponential distribution of the filament length *L* (**fig**. S9**b**). In **fig**. S9**c** we visualise the error landscape by building a PCA from the input vector representing the closeness to the optimal values. We found that many parameter combinations can produce low values of error in eq. (S7), seen in the large cluster of low cost values in **fig**. S9**c I**).

**Table S4:** Treadmilling parameters based on Bayesian optimisation scheme.

| *k*<sub>on</sub> | *k*<sub>off</sub> | *k*<sub>hyd</sub> | *K*<sub>nuc</sub> |
|---|---|---|---|
| 44 s<sup>−1</sup> | 7.03 s<sup>−1</sup> | 34.3 s<sup>−1</sup> | 757 |

### S3.5 Treadmilling parameter scan

**Figure S10:**
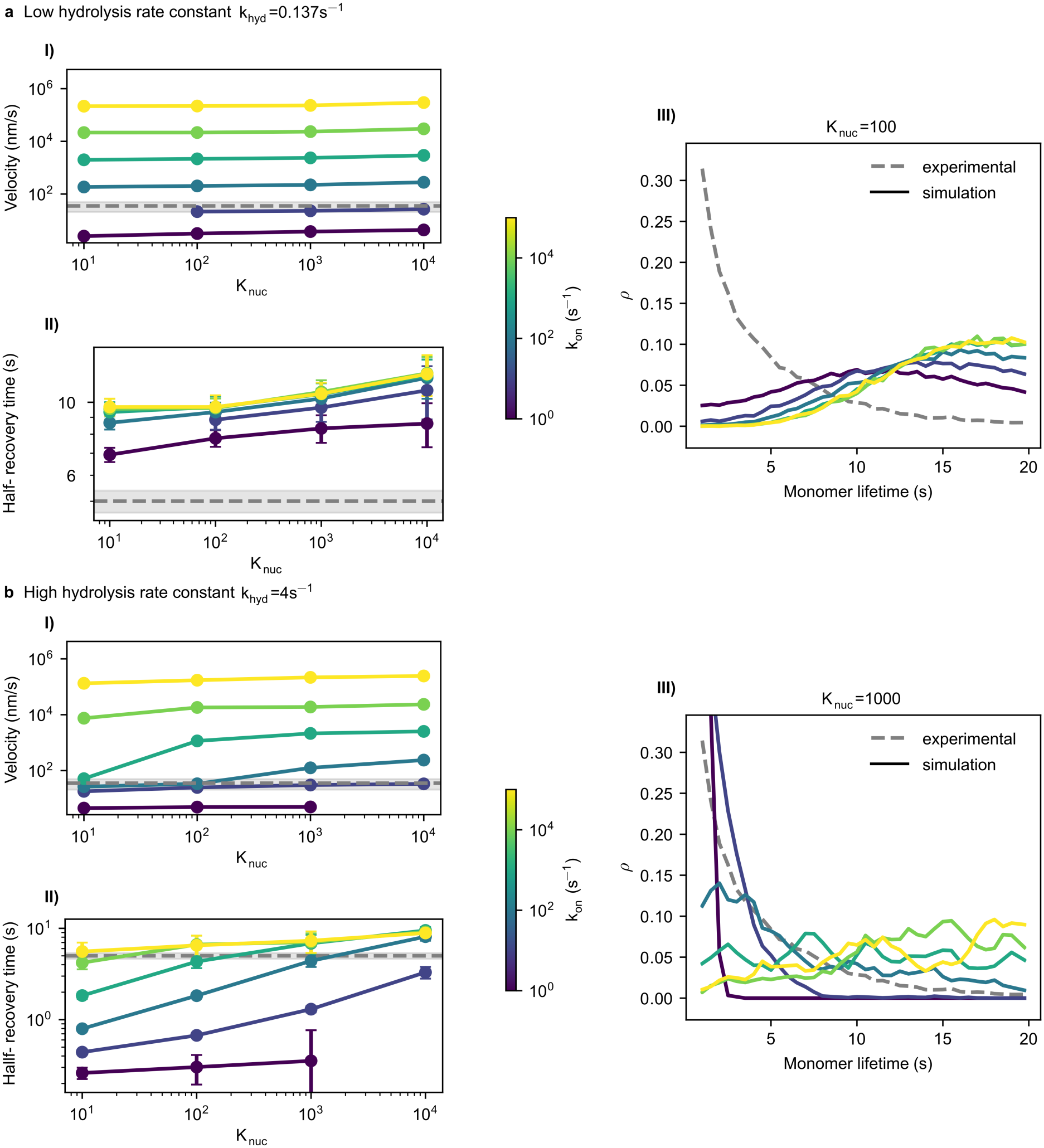
Influence of treadmilling parameters on experimental observables of *in vitro* data [53]. A simulation is run for each combination of *k*_on_ and *K*_nuc_, treadmilling velocity is extracted from ⟨*r*_on_⟩, halfrecovery time from *in silico* FRAP. **a)** Low hydrolysis rate constant *k*_hyd_ based on experimental results [77]. Sample monomer lifetime distributions for *K*_nuc_ = 100 giving a half-recovery time in the right order of magnitude, but Poisson distributed monomer lifetimes. **b)** Hydrolysis rate constant *k*_hyd_ tuned to reproduce exponentially distributed monomer lifetimes, 1-2 orders of magnitude higher than previous assumptions [77, 37]. Sample monomer lifetime distributions given for *K*_nuc_ = 1000, containing the possible best fit to treadmilling velocity, half-recovery time and exponential monomer lifetimes.

